# Brazil Seed Transfer Zones: Supporting Seed Sourcing for Climate-Resilient Ecosystem Restoration

**DOI:** 10.1101/2025.11.20.689464

**Authors:** Mateus Silva, André Braga Junqueira, Pedro Guimarães, Danielle Celentano, Beatriz Murer, Tobias Fremout, Evert Thomas, Christopher Kettle, Edjane Damasceno, Adler Medeiros, Rodrigo Dutra-Silva, Fabian Borghetti, Fatima Piña-Rodrigues, Anabele Gomes, Veronica Maioli, R. Toby Pennington, Lucy Rowland, Eduardo Malta

## Abstract

Ecosystem restoration is key to jointly address the biodiversity, climate, and social inequality crises. In Brazil, large-scale seed-based restoration is gaining momentum, but a national framework to match seed sources with planting sites is still lacking, undermining long-term success.
We developed the first Seed Transfer Zones (STZs) for Brazil using 25 environmental variables. We assessed restoration supply and demand within each STZ and projected the zones into the future under contrasting climate change scenarios.
We identified 48 STZs, distributed across Brazil’s six major vegetation regions (Amazon, Cerrado, Atlantic Forest, Caatinga, Pantanal, and Pampa). Overlaying the STZs with data on seed collection capacity and restoration opportunities revealed priority regions where seed supply is unlikely to meet demand (e.g., Cerrado). Future climate projections also indicate substantial reshaping of STZs, with 51–88% of Brazil’s land area expected to experience mismatches between current and future seed zones.
The creation of Brazil’s first STZs provides a scalable framework to guide seed sourcing for restoration, aligning supply with demand, and future-proofing projects to deliver lasting benefits for biodiversity, climate, and society.

**Societal Impact Statement:** Tens of millions of hectares of degraded land in Brazil are in need of ecosystem restoration to achieve national and global biodiversity and climate goals. Sowing genetically diverse native seeds is key to restoring degraded lands to climate-resilient landscapes, but we still lack guidance on how to effectively match seed sources to restoration sites across broad geographic scales, within the context of a changing climate. We addressed this by identifying 48 Seed Transfer Zones (STZs) based on current and expected future climatic conditions. These zones will help ensure that the right seeds are used in the right places, enabling more effective and climate-smart restoration for nature and people.

## Introduction

Ecosystem restoration has emerged as an integrated solution to simultaneously address biodiversity loss, climate change, and social inequality (Strassburg et al., 2020; Tedesco et al., 2023). The Bonn Challenge aims to restore 350 million hectares of degraded land globally, and the 20x20 Initiative targets 50 million hectares in Latin America by 2030, both aligning with the timeframe of the UN Decade on Ecosystem Restoration (2021–2030) (UNEP, 2021). Within these initiatives, Brazil’s goal is to restore 12 million hectares of native vegetation by 2030, as established under the National Native Vegetation Plan (PLANAVEG) (MMA, 2024). A significant portion of Brazil’s restorable area is unlikely to regenerate naturally due to severe degradation (e.g., eroded pastures) and distance from vegetation remnants, making active restoration critical to trigger ecosystem recovery (Pedrini and Dixon, 2020). Nursery-raised seedling planting has been the most common restoration method, but direct seeding is gaining momentum as it maximises species diversity, reduces implementation costs, and provides income to collectors, many of whom are Indigenous Peoples and Local Communities (IPLCs) (Rodrigues *et al*., 2019; Raupp *et al*., 2020; Ferreira *et al*., 2023). Regardless of the chosen technique, meeting global and regional restoration goals will require unprecedented volumes of native seeds, highlighting the urgent need to strengthen seed supply systems in Brazil and elsewhere (Schmidt *et al*., 2019; Atkinson *et al*., 2021; Bosshard *et al*., 2021; Silva *et al*., 2022; Giacomini *et al*., 2023; Padovezi *et al*., 2024).

Successful restoration in Brazil and globally depends not only on seed quantity but also on seed quality, which is directly associated with its origin (Thomas *et al*., 2014; Erickson and Halford, 2020). Seeds from the same species but different provenances may vary genetically, leading to differences in germination requirements and physiological tolerances (Gallagher and Wagenius, 2016; Aspalter *et al*., 2025). Restoration practitioners traditionally have prioritised locally sourced seeds to reduce the risks of maladaptation, assuming that local provenancing maximises success (Lortie and Hierro, 2022). However, in the context of a changing climate, local seed sources may become maladapted to the future conditions in which restored ecosystems must persist (Broadhurst *et al*., 2008; Breed *et al*., 2013; Havens *et al*., 2015). Practitioners are increasingly following climate-informed seed provenance strategies, collecting seeds primarily from local sources to preserve local adaptation while also including a portion from areas whose current climate resembles the projected future conditions at the restoration site, thereby enhancing adaptability (Prober *et al*., 2015; Ramalho, Byrne and Yates, 2017). Developing seed sourcing strategies in Brazil that explicitly consider future climate scenarios may therefore be essential to support the long-term success of restoration efforts and to enhance the resilience of the country’s ecosystems in a rapidly changing world.

Seed transfer zones (STZs) are a practical tool to guide seed sourcing in ecosystem restoration. STZs represent geographic areas within which seeds can be moved and used with minimal genetic risk, such as maladaptation or outbreeding depression (Hufford and Mazer, 2003). When compared with existing seed sources and restoration needs, STZs also help identify regions where seed supply may lag behind demand, informing strategic investments in seed production systems (De Vitis *et al*., 2017). This can additionally help target the conservation of genetic resources under threat (Gaisberger *et al*., 2022). Ideally, STZs are defined using broad patterns of genetic structure across multiple species (Durka *et al*., 2017; Fremout *et al*., 2021). However, such data are scarce for many regions, especially in the species-rich tropics. For instance, Brazil is home to around 35,000 plant species (Filardi *et al*., 2018), approximately 10% of the global vascular plant flora (Freiberg *et al*., 2020), yet genetic data are available for only a few species (Rull and Carnaval, 2020). In these cases, STZs can be provisionally defined using environmental variables such as climate and soil, which are expected to shape genetic differentiation (Bower, Clair and Erickson, 2014). Because these STZs are based on environmental conditions, they can also be projected into the future using climate models, helping to future-proof restoration projects in the face of changing climates (Marinoni *et al*., 2021).

Despite Brazil’s importance for achieving global restoration goals, the country still lacks a formal STZ system (Dutra-Silva, Overbeck and Müller, 2024). Brazil’s native vegetation is classified into six major regions: the Amazon Rainforest, Cerrado Savanna, Caatinga Dry Forest, Atlantic Rainforest, Pantanal Wetland, and Pampa Subtropical Grassland (IBGE, 2019). Nearly 35% of Brazil’s native vegetation has already been cleared for agriculture (MapBiomas, 2025). Under Law 12.651 of 2012, rural landowners must retain native vegetation on at least 20% of their land, increasing to 80% in the Amazon. Landholdings below this threshold are legally required to restore degraded areas, creating a national restoration deficit of 19.3 million hectares in 2022 (OCF, 2025), 7.3 million hectares above the 2030 target set by PLANAVEG. Additionally, pastures represented 56% of Brazil’s land under anthropic use in 2023, and 22% of these pastures face severe degradation (MapBiomas, 2025), representing a substantial demand for restoration outside legal obligations (Barros *et al*., 2023; Lewis *et al*., 2023). In response, a growing number of seed suppliers have emerged across the country, many organised as cooperatives and associations of IPLCs who collect, process, and sell native seeds (Urzedo *et al*., 2022). More recently, these initiatives have been coordinated through the Redário Coalition, which centralises seed orders and redistributes them to local suppliers (Celentano *et al*., 2024). The coalition brings together more than 30 seed networks across Brazil. Beyond coordinating seed trading, Redário also provides training and technical support, expands market access, and fosters cooperation between suppliers and upstream actors in the seed supply chain. Currently, Redário limits seed exchange to Brazil’s six major vegetation regions, without incorporating climate change considerations into seed provenance decision-making. Developing STZs for Brazil will equip Redário and other stakeholders with a science-based framework to match seed sources with restoration sites, enabling more effective and resilient restoration while contributing to biodiversity conservation, climate mitigation, and social equity.

In this study, we aimed to (1) develop provisional Seed Transfer Zones (STZs) for Brazil, (2) characterise these zones in terms of their restoration status, and (3) project them into future climate scenarios. To delineate the zones, we performed a clustering analysis using 25 climatic and edaphic variables. We then characterised each zone using data on Redário seed collector density, recent land use and land cover, the legal restoration deficit, and the extent of severely degraded pastures. Finally, we projected the zones to the middle and end of the 21^st^ century using nine climate forecasts under both low- and high-emission scenarios. All R code for the analysis, as well as a Shiny App for interactive online visualisation of the STZs, are openly available at https://github.com/silva-mc/STZ-BR. These findings advance ecosystem restoration in Brazil while promoting biodiversity conservation and socio-economic benefits.

## Material and methods

### Environmental data

We downloaded 19 bioclimatic variables (BIO1–BIO19) from WorldClim version 2.1 for the 1970–2000 period using the geodata R package (Hijmans *et al*., 2024). To avoid artefacts that generate artificial discontinuities in Brazil, we excluded precipitation of the warmest (BIO18) and coldest quarters (BIO19) from subsequent analyses (Booth, 2022). We also retrieved eight edaphic variables from SoilGrids version 2.0 at a 5–15 cm depth, a common range for nutrient analyses, using geodata (Poggio *et al*., 2021). All layers were cropped and masked to the mainland extent of Brazil, yielding a harmonised raster stack with 25 layers (17 climatic and 8 edaphic variables) at 150 arcsec resolution (∼5 km at the Equator; Figure S1).

We performed a Principal Component Analysis (PCA) on the climatic and edaphic variables to reduce dimensionality and address collinearity. The raster stack was first converted into a data frame, and cells with missing values (e.g., water bodies) were removed. The PCA retained the first five principal components (PCs) using eigenvalues > 1 as the criterion (ranging from 1.15 to 9.89, cumulative variance of 86.13%). To enhance the spatial structure of the resulting clusters, we combined PC scores with scaled latitude and longitude per pixel. Finally, we converted the PC scores and coordinates back to raster format, producing a final environmental dataset composed of seven variables (Table S1, Table S2, Figure S2).

### Cluster analysis

We conducted cluster analysis on the seven environmental variables using the CLARA algorithm implemented in the cluster R package (Maechler et al., 2023). CLARA is designed for large datasets and is more robust to outliers than k-means. Briefly, it repeatedly samples subsets of the data, applies Partitioning Around Medoids (PAM) to each, and selects the clustering with the lowest average dissimilarity across the full dataset. Each observation is then assigned to its nearest medoid.

Because cluster analysis requires defining the number of clusters (k) a priori, we ran the CLARA algorithm 50 times, testing k values from 20 to 70. We used Euclidean distance to construct the dissimilarity matrix and ran 50 samples of 2,000 observations each. After each run, we projected the clusters in geographical space, and values were reassigned using a 3 × 3 moving window (centred on each grid cell), based on the most frequent cluster among the 8 neighbouring cells, thereby reclassifying isolated pixels and improving spatial consistency.

We selected the final STZ map as the clustering version with the lowest k for which the mean range in annual temperature (BIO1) within clusters was below 3 °C (Figure S3). To determine this, we stacked each of the 50 STZ versions, converted them to a data frame, and calculated the 2.5^th^ and 97.5^th^ percentiles of BIO1 within each cluster. We then computed the 95% range per cluster, averaged the BIO1 range across clusters for each STZ version (and its corresponding k), and plotted the mean BIO1 range against k. This criterion was adapted from the U.S. STZ framework (Bower, Clair and Erickson, 2014), which is based on empirical evidence of maladaptation when seed transfers exceed 3 °C between donor and recipient sites (St. Clair, Howe and Kling, 2020).

To assess environmental similarities among clusters, we performed hierarchical clustering. For each cluster, we first calculated the mean of the seven variables used to delimit the STZs under present conditions. We then computed a Euclidean distance matrix among clusters and applied a modified Ward’s minimum variance method using the square root of dissimilarities. The resulting hierarchical object was converted into a dendrogram.

### Restoration profile

We used the density of Redário seed collectors per zone as a proxy for seed supply capacity for ecological restoration. We first identified 31 cooperatives, associations, and other organisations affiliated with the Redário Coalition (Table S3). Because the exact location of the seed collection areas was not available for most suppliers, we approximated seed collection sites using the centroids of the municipalities in which each supplier operates. We then obtained the number of seed collectors per supplier from Redário’s 2024 census and divided this number by the number of municipalities per supplier to estimate the number of collectors per seed collection site (municipality). For four suppliers not included in the census, we assumed a single seed collector each. Finally, we calculated the number of seed collectors per STZ by extracting the most common STZ within a 5 km buffer around each municipality, summing the number of collectors per STZ, and dividing this value by the STZ area, standardised to a spatial scale of 100,000 km^2^.

We used native vegetation cover as an additional proxy for seed supply capacity because native vegetation remnants are the main source of seeds for restoration in Brazil (Schmidt *et al*., 2019). To estimate the native vegetation cover of each STZ, we retrieved the land-use and land-cover (LULC) data of 2023 from MapBiomas Collection 9 (MapBiomas, 2025). We extracted only natural vegetation, specifically: forest, savanna, mangrove, wetland, grassland, rock outcrop, and coastal vegetation. The LULC layer, with a resolution of ∼1 arcsec, was aggregated to match the lower resolution of the STZ raster (150 arcsec). Each aggregated grid cell value represented the fraction of native vegetation within that area. This fraction was then multiplied by the cell area, summed across each STZ, and divided by the total STZ area to calculate the percentage of native vegetation per STZ.

We used the legal restoration deficit as an indicator of restoration demand across Brazil. The legal restoration deficit is based on the latest revision of rural property compliance with Law 12.651 of 2012 (commonly known as the Forest Code), available from the *Termômetro do Código Florestal* platform (https://termometroflorestal.org.br/; OCF, 2025). The data indicate the area within each Brazilian municipality that is legally required to undergo ecological restoration, either in permanent preservation areas or legal reserves. We acknowledge that the Forest Code permits landowners to restore their lands through multiple approaches, including the use of a set percentage of exotic species; however, we assume that native seeds will still be required for restoration implementation in most cases (e.g., native trees in agroforestry). For each municipality, we calculated the proportion of its area falling within each STZ and multiplied the municipality’s restoration deficit by this proportion, resulting in the area under restoration deficit per STZ per municipality. We then summed the restoration deficit areas across municipalities within each STZ and divided by the STZ total area to obtain the percentage of the STZ that requires ecological restoration for legal compliance.

We used degraded pasture cover as an additional metric of seed demand for restoration. Similarly to the LULC routine, we used 2023 pasture quality data from MapBiomas Collection 9 (MapBiomas, 2025), selected pixels with the lowest quality score (1), aggregated them to the 150 arcsecond STZ resolution, and calculated the percentage of degraded pasture per STZ.

### Future projections

We retrieved bioclimatic variables BIO1 to BIO17 for the 2041–2060 and 2081–2100 periods under two Shared Socioeconomic Pathways (SSP1 and SSP5), using nine climate models from CMIP6 at 150 arcsec resolution via the geodata R package (Figures S4–S7) (Eyring *et al*., 2016; Hijmans *et al*., 2024). SSP1 (“Sustainability”) represents an optimistic scenario with mitigated emissions and global warming limited to 2 °C, while SSP5 (“Fossil-fuelled Development”) reflects a high-emission scenario with projected warming above 4 °C by the end of the 21^st^ century.

We initially selected five high-performance models at the global scale (Brunner *et al*., 2020) and another five at the Brazil scale (Reboita *et al*., 2024). Because one model (MPI-ESM1-2-HR) was common to both sets, the final list comprised nine models: GISS-E2-1-G, FIO-ESM-2-0, ACCESS-CM2, AWI-CM-1-1-MR, EC-Earth3-Veg, INM-CM4-8, IPSL-CM6A-LR, CMCC-ESM2, and MPI-ESM1-2-HR. Edaphic and topographic variables were not modified.

Using the PCA object derived from the present data, we projected the 25-layer future environmental dataset onto the same principal components for each SSP and timeframe, processing one model at a time. We retained the first five PCs and added the rescaled latitude and longitude, resulting in seven future variables. Each future observation was then assigned to the nearest existing medoid from the CLARA clustering object that generated the final STZ map, using Euclidean distance for each model. The same spatial smoothing, resampling, and masking routines were applied. The final outputs consisted of four maps of projected STZ distributions for 2060 and 2100 under SSP1 and SSP5 for each of the nine climate models.

We calculated two indices to describe the future distribution of STZs across the nine climate models. The first was an uncertainty index, defined as the number of different STZs predicted by the models for a given timeframe and scenario. An uncertainty value of one indicates complete agreement among models, whereas a value of nine indicates that each model projects a different future STZ. The second was a mismatch index, defined as the number of models in which the future STZ differs from the current STZ. A mismatch of zero means all models project the same STZ as the present, while a mismatch of nine means all models agree that the future STZ differs from the current one. We developed an R-based Shiny App providing practitioners with interactive visualisation of STZs across current and future timeframes (https://mateus-silva.shinyapps.io/stz-br/). All analyses were performed on the R environment version 4.4.1 (R Core Team, 2024).

## Results

### Brazil Seed Transfer Zones (STZs)

We identified 48 Seed Transfer Zones (STZs) across Brazil’s six major vegetation regions, ranging in size from 35,080.55 (STZ 44) to 369,476.64 km^2^ (STZ 3; Figure 1), corresponding to 0.42% to 4.42% of Brazil’s area, respectively. The absolute number of STZs per biome varied from five (Pantanal) to 32 (Amazon). Excluding STZs that represented less than 5% of the biome area reduced the number of STZs per biome to two in the Pantanal, three in the Pampa, seven in both the Amazon and Caatinga, nine in the Atlantic Forest, and 11 in the Cerrado. Environmental similarities among these STZs are shown in Figure S8.

**Figure 1.**
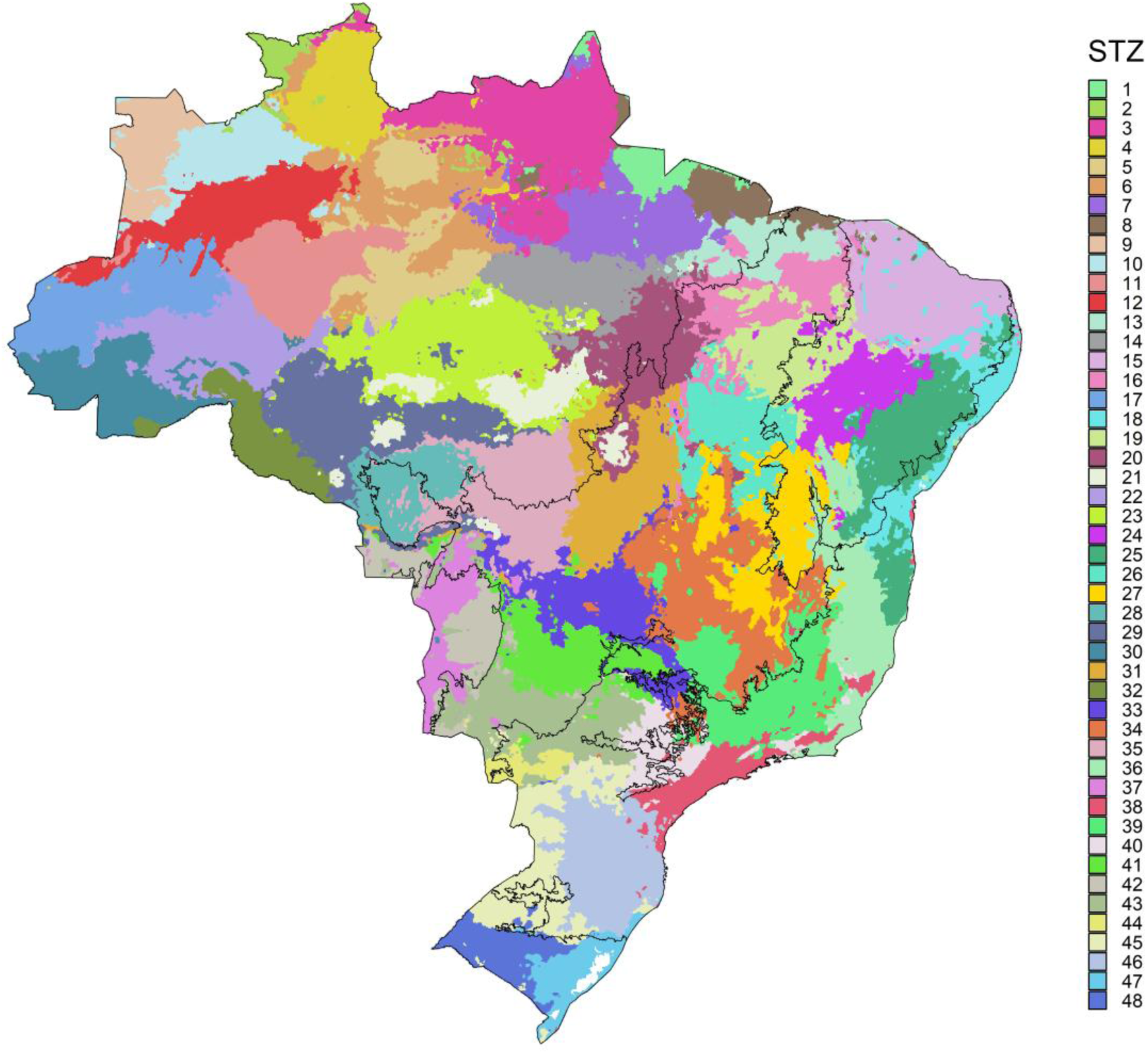
Provisional Seed Transfer Zones (STZs) for Brazil. Each colour represents one of 48 STZs defined by a clustering analysis based on climatic and edaphic variables. Black contours delineate Brazil’s six major vegetation regions (Amazon, Cerrado, Caatinga, Atlantic Forest, Pantanal, and Pampa).

### Seed supply and demand per STZ

As of June 2025, the *Redário* coalition had 32 active seed suppliers operating across 176 municipalities and employing ∼2,578 seed collectors throughout Brazil. The estimated number of seed collectors per zone ranged from zero (in 21 STZs) to 643 in STZ 29 (southern Amazon). After excluding STZs with no seed collection activity and standardising by area, collector density ranged from 0.15 (STZ 41) to 181.5 per 100,000 km^2^ (STZ 29), with an average of 34.7 collectors per 100,000 km^2^ (Figure 2a).

**Figure 2.**
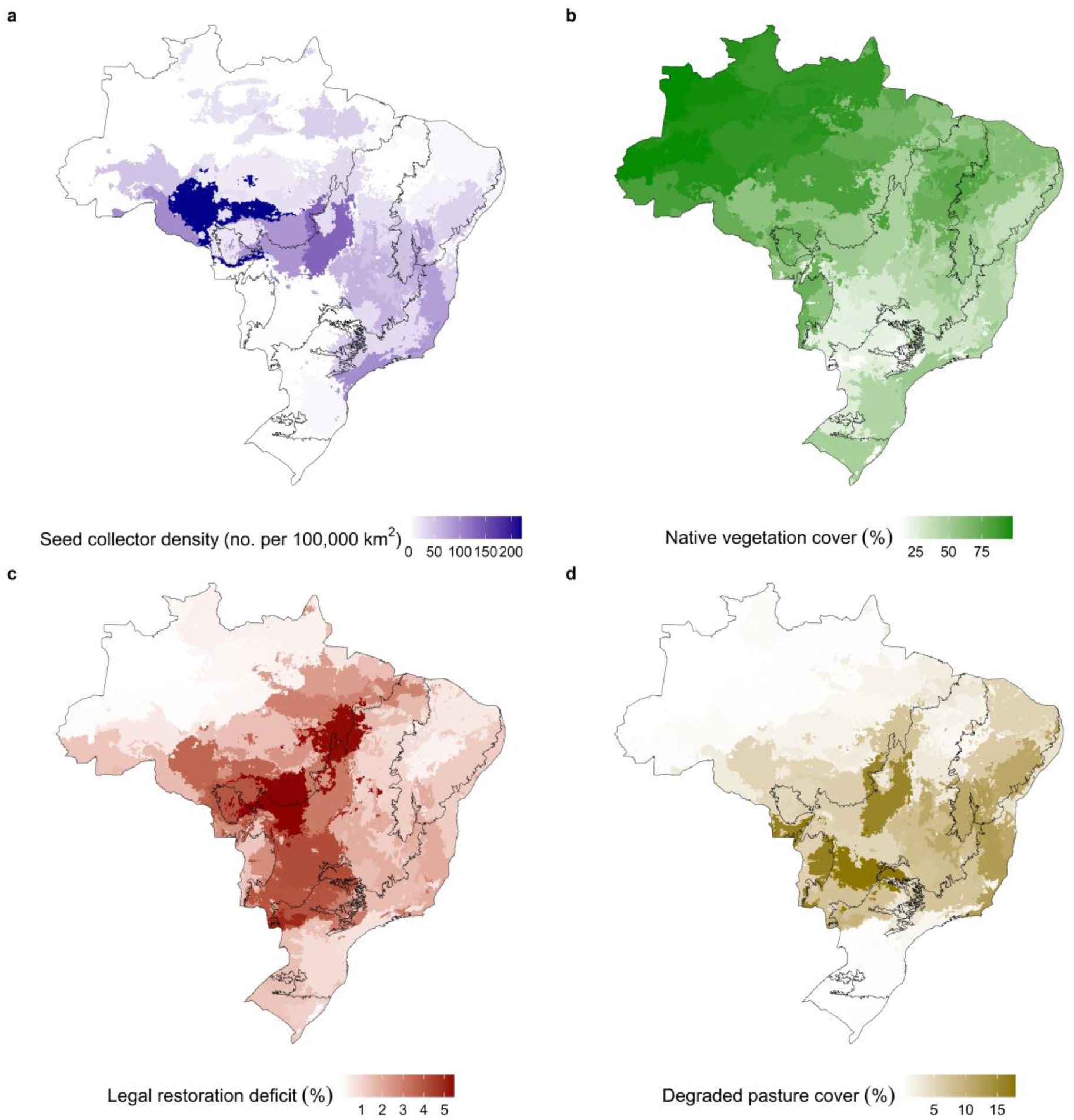
Seed supply and demand across Brazil’s Seed Transfer Zones (STZs). Seed supply was represented by (**a**) the density of seed collectors affiliated with the Redário Coalition and (**b**) the extent of native vegetation cover in 2023 per STZ. Seed demand was represented by (**c**) the legal restoration deficit under the Forest Code and (**d**) the extent of degraded pasture cover in 2023 per STZ. Black contours delineate Brazil’s six major vegetation regions.

STZ 3 (northern Amazon) was not only the largest in total area but also the zone with the greatest extent of conserved native vegetation in 2023, totalling 330,900.59 km^2^ (Figure 2b). When considering the proportion of native vegetation relative to total area, STZ 9 in the western Amazon had the highest cover, with 98.39% of its area remaining native in 2023. In contrast, STZ 44 in the Cerrado–Atlantic Forest transition had the lowest coverage of native vegetation in both absolute terms (4,481.29 km^2^) and relative terms (12.83%).

In 2022, Brazil had 193,320 km^2^ of rural private land classified as being in legal restoration deficit, according to Law 12.651 of 2012. STZ 35 in the Amazon–Cerrado transition had the largest deficit expressed as absolute area (15,696.07 km^2^) and relative to the zone’s area (5.46%; Figure 2c). Based on relative deficit, STZ 20 ranked second (5.37%), STZ 44 third (4.95%), STZ 33 fourth (4.16%), and STZ 41 fifth (4.15%), all located in the Cerrado.

Pastures classified as severely degraded covered 362,181.99 km^2^ of Brazil, representing approximately 4.26% of the national territory. STZ 31 in central-western Cerrado had the highest absolute degraded pasture cover (34,559.06 km^2^), while STZ 41 in southern Cerrado had the highest relative cover (17.88%; Figure 2d). Following STZ 41, STZs 42 (eastern Pantanal), 31, 36 (central Atlantic Forest), and 25 (south-eastern Caatinga) also showed high degraded pasture coverage, ranging from 11 to 16%.

### Future STZs

When projecting the STZs to the 2041–2060 timeframe and using the MPI-ESM1.2-HR climate model, STZ 18 was the zone with the largest contraction relative to its current extent under SSP1 scenario (−59.90%) and STZ 28 under the SSP5 scenario (−92.96%; Figure 3). As for the 2081–2100 timeframe, STZ 14 showed reductions of −61.02% under SSP1 and STZ 28 found no climate analogue under SSP5, resulting in an area loss of 100%. STZ 13 showed the largest expansion in both timeframes and scenarios (2060 SSP1: 87.37%, 2060 SSP5: 252.92%, 2100 SSP1: 99.87%, 2100 SSP5: 1151.99%).

**Figure 3.**
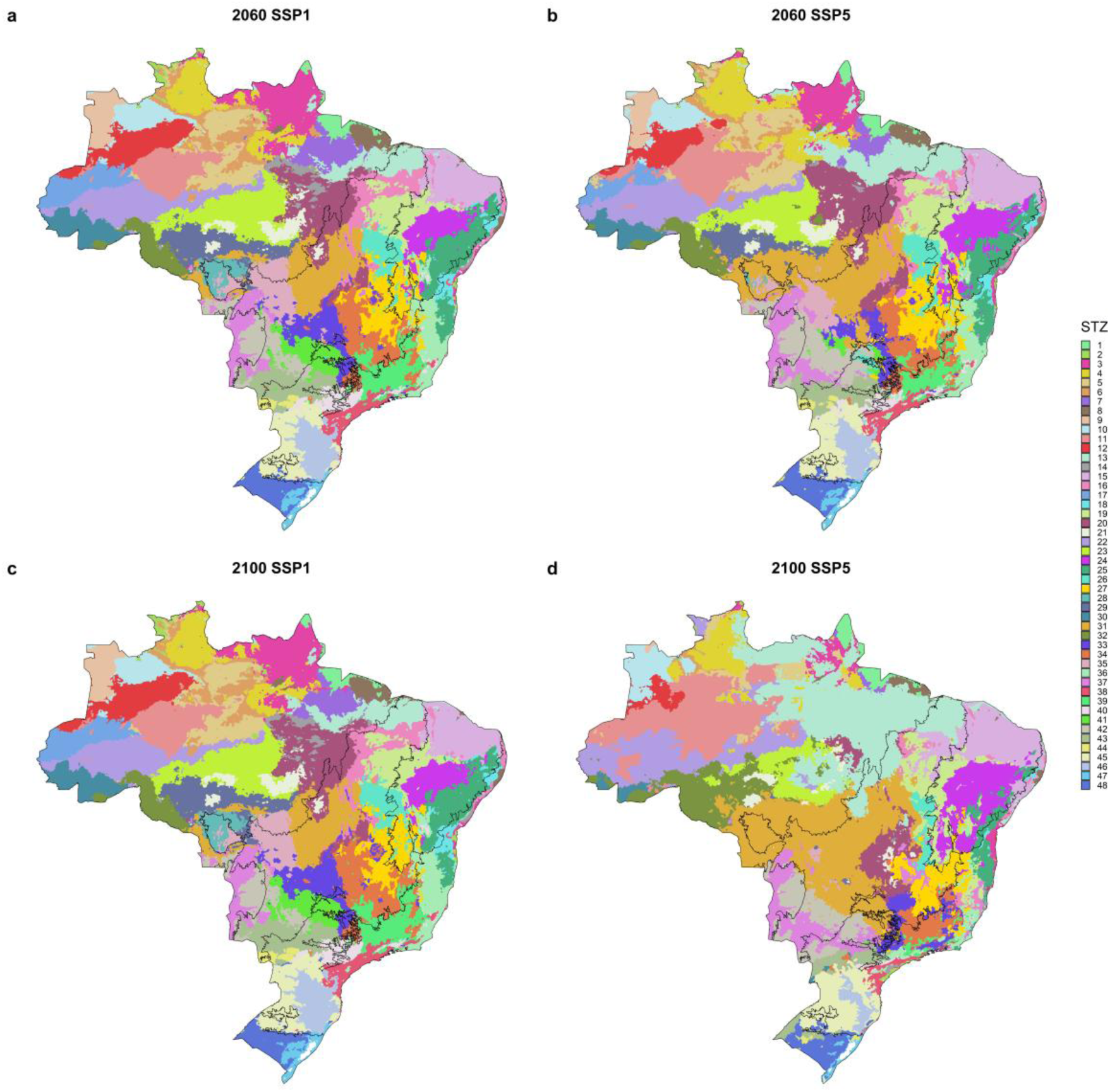
Projected Seed Transfer Zones (STZs) under future climatic scenarios. Projections are based on the MPI-ESM1.2-HR climate model for (**a**, **b**) 2041–2060 and (**c**, **d**) 2081–2100 under (**a**, **c**) low (SSP1) and (**b**, **d**) high (SSP5) greenhouse gas emission scenarios. Black contours delineate Brazil’s six major vegetation regions.

STZ predicted for the future varied across climate models. Between 40.03% (SSP1) and 51.17% (SSP5) of the national territory had two or more climate models predicting different STZs by 2060 (uncertainty > 1; Figure 4). This area of disagreement increased toward 2100, reaching 48.76% (SSP1) and 72.66% (SSP5) of the country. The maximum uncertainty was six different STZs predicted for the same location, but occurring in only 0.19% of Brazil by 2100 under SSP5 and in less than 0.01% under the other timeframes and scenarios.

**Figure 4.**
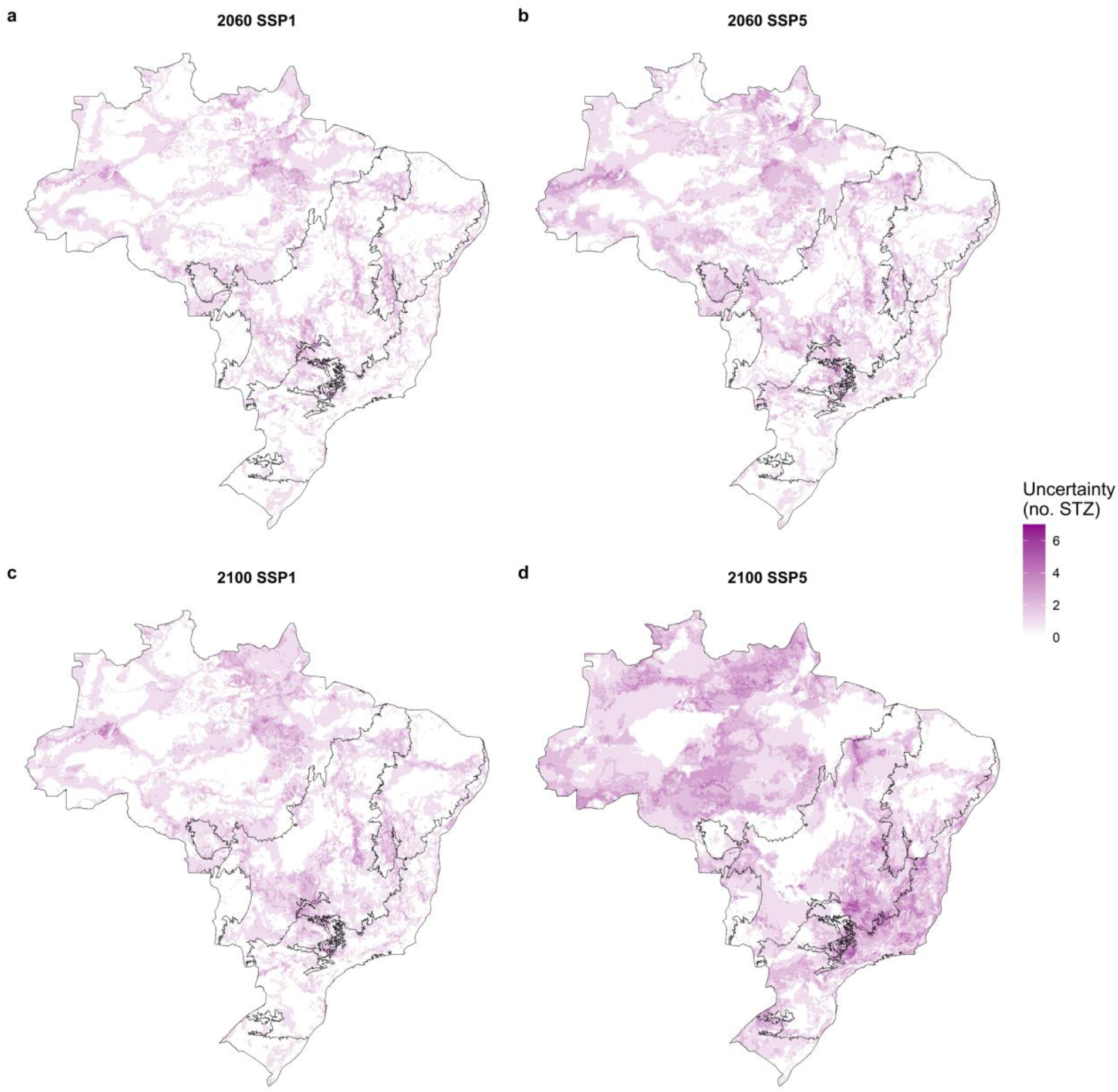
Uncertainty in future projections of Brazil’s Seed Transfer Zones (STZs). Uncertainty represents the number of distinct STZs predicted by nine climate models, estimated for (**a**, **b**) 2041–2060 and (**c**, **d**) 2081–2100 under (**a**, **c**) low (SSP1) and (**b**, **d**) high (SSP5) greenhouse gas emission scenarios. Black contours delineate Brazil’s six major vegetation regions.

By 2060, 51.32% (SSP1) and 66.28% (SSP5) of Brazil are projected to have climate conditions that more closely resemble those of a different STZ than the current one (mismatch > 0; Figure 5). Considering 2100, the area expected to experience a mismatch between present and future STZs increased to 57.76% under SSP1 and 88.44% under SSP5. The maximum mismatch level (9) extended across 18.12% (2060 SSP1), 27.90% (2060 SSP5), 16.45% (2100 SSP1), and 54.08% (2100 SSP5) of Brazil’s territory.

**Figure 5.**
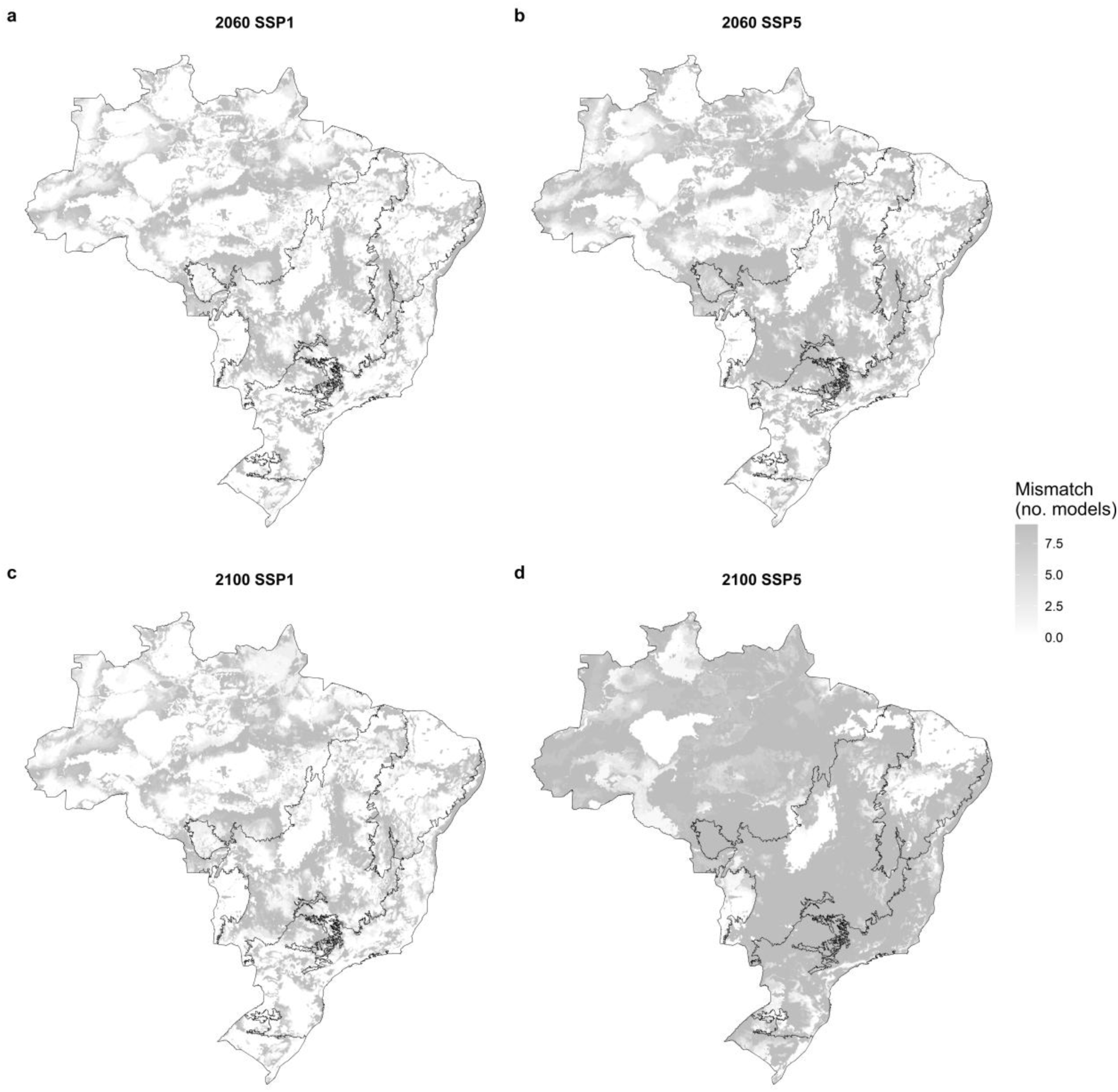
Mismatch between current and future Seed Transfer Zones (STZs). Mismatch represents the number of climate models predicting a future STZ that differs from the current one, calculated for (**a**, **b**) 2041–2060 and (**c**, **d**) 2081–2100 under (**a**, **c**) low (SSP1) and (**b**, **d**) high (SSP5) greenhouse gas emission scenarios. Black contours delineate Brazil’s six major vegetation regions.

## Discussion

In this study, we delineated 48 Seed Transfer Zones (STZs) to guide seed sourcing for ecosystem restoration in Brazil. These zones vary in size, spanning approximately 0.4% to 4% of the country’s territory. STZs in the Amazon showed the highest levels of seed supply capacity based on the density of Redário seed collectors and the extent of native vegetation cover. In contrast, STZs in the Cerrado concentrated the greatest restoration opportunities, as indicated by legal restoration deficit and degraded pasture cover. Climate change is projected to shift the distribution of STZs by mid- and late-century. Although climate models differ substantially in their forecasts, up to 88% of Brazil’s territory could experience climates that no longer match the envelope of current zones by the end of the 21^st^ century under a high-emissions scenario. These findings call for strategic, climate-informed seed procurement to support Brazil’s ambitious restoration goals in the decades ahead.

The 48 proposed STZs advance the seed sourcing guidelines for Brazil. Redário is the main organisation coordinating the seed supply chain in Brazil, and its current provenancing principle aims to restrict seed movement between donor and restoration sites by using the boundaries of Brazil’s six major vegetation regions as defined by IBGE, which are based on floristic similarity as well as climate, soil, and topography (IBGE, 2019). Our STZs provide a more nuanced framework, moving from six to 48 spatial units that often span multiple regions and capture transition areas between IBGE’s classifications. Previous studies performed similar analysis for specific regions, such as the Cerrado (Sano *et al*., 2019; Cattelan *et al*., 2024) and Pampa (Hasenack *et al*., 2023), but ours was the first to apply the STZ framework nationwide, matching the scale at which Redário operates. However, organisations working at finer spatial scales (e.g., watershed level) may require alternative STZ delineations with more and smaller zones (Ximenes *et al*., 2021). To support this flexibility, we provide open-source code to replicate our analyses with any number of STZs (https://github.com/silva-mc/STZ-BR). While finer-scale zoning may be desirable in some contexts, there are trade-offs. A large number of zones, as seen in Germany, which has 22 STZs, further divided into 72 subzones (Durka *et al*., 2025), can complicate restoration logistics by reducing seed availability within zones (Mainz and Wieden, 2019). For Brazil’s scale, the scope of this study, we argue that 48 STZs strike a practical balance, enabling suppliers to secure both the quantity and quality of seeds needed for ambitious restoration efforts across the country.

Our STZ map was designed to match restoration sites with suitable seed sources, but an alternative use is identifying zones where seed demand outpaces supply capacity. For instance, STZs 35 and 20, along the Amazon Arc of Deforestation, have a high legal restoration deficit, likely resulting from rapid land-use change and from strict conservation requirements that mandate up to 80% of rural properties in the Amazon remain under natural vegetation (Camara *et al*., 2023). Yet, the highest density of seed collectors is in STZ 29, which borders STZs 35 but extends rather into the interior of the Amazon. The concentration of seed collectors in STZ 29 is largely explained by the presence of three major seed suppliers: the Xingu Seed Network Association, Brazil’s oldest and largest native seed supplier (Campos-Filho *et al*., 2013; Durigan, Guerin and da Costa, 2013), the Amazon Portal Seed Network, and the Amazonian Bioeconomy Seed Network. Together, these networks involve more than 1,400 collectors (ARSX, 2025; RESEBA, 2025; RSPA, 2025). STZs 33 and 41, in the southern Cerrado, face a similar gap because they have the lowest density of seed collectors yet rank among the zones with the largest areas requiring legal restoration and with extensive severely degraded pastures. By revealing where restoration demand is high but supply is lacking, our STZ framework offers a practical tool to prioritise seed supply development. This includes training seed collectors, improving storage infrastructure, and strengthening local seed governance to support Brazil’s restoration ambitions (Urzedo *et al*., 2022; Padovezi *et al*., 2024).

Climate change will drive a significant redistribution of STZs across Brazil. More than half of the country is projected to experience a mismatch between current and future STZs by the end of the 21^st^ century under a high-emissions scenario (SSP5). In such high-risk areas, practitioners may adopt a climate-informed provenancing strategy by sourcing seeds from both present and projected STZs (Prober *et al*., 2015). Yet, the future STZ per locality was highly dependent on the climate model used and the emissions scenario. To account for such uncertainties, we proposed an adaptation of the “portfolio” approach (Fremout *et al*., 2021), where: 50% sourced from within the current STZ, 20% from the STZs forecasted under a low-emission scenario (SSP1), and 30% under a high-emission scenario (SSP5). The 50% of seeds matching the future STZ would be divided equally across the nine climate models. Practitioners could also consider the 2060 timeframe for projects using short-lived species (e.g., grasses, forbs, shrubs) and 2100 for long-lived species (e.g., trees). This approach further highlights opportunities to expand seed supply networks in climatically strategic STZs. For example, STZ 13, in the northern Cerrado, is expected to expand substantially across all timeframes and scenarios, yet currently has limited representation in the seed supply chain. Without action, this gap may hinder the inclusion of adaptive genetic material in seed mixes. Since long-distance seed exchange is already a common practice in Brazil (Dutra-Silva, Overbeck and Müller, 2024), incorporating future STZ projections into seed planning can help guide seed movement to areas where it strengthens the climate resilience of restored populations and improves the likelihood of long-term restoration success.

We highlight five key considerations for the application of STZs in Brazil and elsewhere. First, while STZs provide a practical framework for most species, they are not universally applicable. Range-restricted or threatened species may occur entirely within a single STZ, requiring either finer delimitation or the use of more targeted tools such as the Climate-Oriented Seed Sourcing Tool (COSST) (Silva *et al*., 2025). Second, gridded soil products are useful for incorporating edaphic variables known to shape species’ genetic differentiation, but such products carry high uncertainty (Radočaj *et al*., 2023; Lilburne *et al*., 2024) and often lack key variables, such as water table depth. Third, the effective implementation of STZs depends on improving seed traceability. The Redário is currently training collectors to record the precise geographic origin of each seed batch using a mobile app (personal communication); however, traceability remains coarse when seeds from multiple locations, potentially spanning different STZs, are mixed during processing and storage. Addressing this will require improved policies, infrastructure, and protocols. Fourth, empirical validation and ongoing monitoring are essential to refine and support STZ use. Broad-scale genetic studies across species can help optimise STZ delineation by identifying consistent genetic clustering patterns (Jørgensen *et al*., 2016; Durka *et al*., 2017), while common garden experiments provide further validation by testing whether seeds from different STZs exhibit distinct trait performance related to growth, survival, or reproduction (Nolan *et al*., 2023; Aoyama *et al*., 2025). Fifth, monitoring restoration outcomes offers real-world feedback, revealing how the composition of seed mixes, specifically the STZs they represent, influences ecosystem trajectories under a changing climate (Thomas *et al*., 2014; Pizza, Foster and Brudvig, 2023). While continuous refinement is both necessary and expected, Brazil’s first STZs provide a strong foundation for guiding seed sourcing for ecosystem restoration across one of the world’s most biodiverse and rapidly changing countries.

## Acknowledgements

LR acknowledges NERC for the Independent Research Fellowship NE/N014022/1 and NERC-FAPESP grant 19/07773-1.

## Author contribution

MS, EM, ABJ, PG, DC, and BM conceptualized the research; MS led the data analysis and drafted the manuscript; all authors contributed to the manuscript during the revision phase.

## Data availability statement

The core data used to reproduce the analysis, together with the R code and the raster file of the STZs, are available in the GitHub (https://github.com/silva-mc/STZ-BR) and Figshare repositories (https://doi.org/10.6084/m9.figshare.30294655, https://doi.org/10.6084/m9.figshare.30294589, https://doi.org/10.6084/m9.figshare.30294523).

