## Supplementary material for "Brazil Seed Transfer Zones: Supporting Seed Sourcing for Climate-Resilient Ecosystem Restoration"

**Supplementary material to the article "Brazil Seed Transfer Zones: Supporting Seed Sourcing for Climate-Resilient Ecosystem Restoration" by Silva et al.**

**Table S1.** Loadings from the Principal Component Analysis (PCA) based on 25 climatic and edaphic variables. The magnitude of each loading indicates the importance of the variable for that principal component, while the sign reflects the direction of the association. BIO1 stands for annual mean temperature (°C); BIO2, mean diurnal temperature range (°C); BIO3, isothermality (%); BIO4, temperature seasonality (standard deviation ×100, °C); BIO5, maximum temperature of the warmest month (°C); BIO6, minimum temperature of the coldest month (°C); BIO7, temperature annual range (°C); BIO8, mean temperature of the wettest quarter (°C); BIO9, mean temperature of the driest quarter (°C); BIO10, mean temperature of the warmest quarter (°C); BIO11, mean temperature of the coldest quarter (°C); BIO12, annual precipitation (mm); BIO13, precipitation of the wettest month (mm); BIO14, precipitation of the driest month (mm); BIO15, precipitation seasonality (coefficient of variation, %); BIO16, precipitation of the wettest quarter (mm); and BIO17, precipitation of the driest quarter (mm). Bulk density is expressed in $kg/dm^3$, coarse fragments in %, nitrogen content in g/kg, pH is unitless, sand, silt, and clay in %, and soil organic carbon (SOC) in g/kg.

| | PC1 | PC2 | PC3 | PC4 | PC5 | PC6 | PC7 | PC8 | PC9 | PC10 | PC11 | PC12 | PC13 | PC14 | PC15 | PC16 | PC17 | PC18 | PC19 | PC20 | PC21 | PC22 | PC23 | PC24 | PC25 |
|---|---|---|---|---|---|---|---|---|---|---|---|---|---|---|---|---|---|---|---|---|---|---|---|---|---|
| BIO1 | 0.29 | -0.15 | -0.04 | 0.05 | -0.05 | 0.11 | 0.03 | -0.10 | 0.01 | 0.01 | 0.01 | -0.03 | -0.05 | 0.04 | 0.05 | -0.08 | 0.11 | 0.45 | -0.04 | 0.11 | -0.27 | 0.71 | 0.20 | 0.00 | 0.00 |
| BIO2 | -0.16 | -0.18 | 0.40 | 0.18 | -0.08 | -0.01 | -0.02 | -0.05 | -0.52 | 0.26 | -0.11 | 0.00 | 0.05 | -0.02 | 0.19 | 0.09 | 0.03 | 0.20 | 0.07 | -0.17 | -0.46 | -0.24 | 0.01 | 0.00 | 0.00 |
| BIO3 | 0.27 | -0.02 | -0.09 | -0.08 | 0.05 | -0.40 | -0.18 | -0.16 | -0.39 | 0.14 | -0.07 | 0.04 | -0.03 | 0.15 | 0.52 | -0.13 | 0.05 | -0.25 | -0.04 | 0.15 | 0.32 | 0.13 | -0.01 | 0.00 | 0.00 |
| BIO4 | -0.28 | 0.09 | -0.09 | 0.14 | 0.00 | 0.16 | 0.31 | 0.06 | 0.06 | -0.33 | -0.09 | -0.01 | -0.10 | 0.34 | 0.42 | 0.07 | 0.13 | -0.20 | 0.03 | 0.03 | -0.24 | 0.18 | -0.43 | 0.00 | 0.00 |
| BIO5 | 0.19 | -0.23 | 0.10 | 0.18 | -0.13 | 0.40 | 0.21 | 0.08 | -0.13 | 0.22 | 0.09 | -0.05 | 0.01 | -0.05 | -0.13 | -0.30 | -0.18 | -0.53 | 0.00 | 0.06 | -0.01 | 0.12 | 0.00 | 0.00 | 0.36 |
| BIO6 | 0.30 | -0.05 | -0.19 | -0.07 | -0.01 | 0.02 | -0.01 | -0.03 | 0.08 | 0.05 | 0.08 | 0.00 | -0.04 | -0.02 | -0.01 | -0.20 | -0.08 | -0.28 | 0.07 | -0.10 | -0.41 | -0.12 | 0.00 | 0.00 | -0.73 |
| BIO7 | -0.26 | -0.08 | 0.29 | 0.19 | -0.07 | 0.21 | 0.14 | 0.09 | -0.17 | 0.07 | -0.04 | -0.04 | 0.05 | -0.01 | -0.07 | 0.07 | -0.01 | 0.03 | -0.09 | 0.16 | 0.50 | 0.22 | -0.01 | 0.00 | -0.59 |
| BIO8 | 0.23 | -0.16 | -0.01 | 0.10 | -0.07 | 0.08 | 0.02 | -0.46 | 0.01 | -0.28 | -0.62 | 0.08 | 0.19 | -0.12 | -0.13 | 0.34 | -0.06 | -0.16 | -0.01 | 0.00 | 0.02 | -0.02 | 0.00 | 0.00 | 0.00 |
| BIO9 | 0.29 | -0.10 | -0.10 | -0.01 | -0.01 | 0.12 | 0.06 | 0.13 | 0.01 | 0.19 | 0.35 | -0.05 | -0.16 | 0.00 | 0.13 | 0.81 | -0.01 | -0.08 | -0.01 | -0.01 | 0.03 | -0.02 | 0.00 | 0.00 | 0.00 |
| BIO10 | 0.25 | -0.15 | -0.13 | 0.14 | -0.07 | 0.31 | 0.27 | -0.07 | 0.05 | -0.11 | -0.01 | -0.04 | -0.08 | 0.28 | 0.24 | -0.20 | 0.18 | 0.31 | 0.03 | -0.07 | 0.26 | -0.47 | 0.27 | 0.00 | 0.00 |
| BIO11 | 0.29 | -0.13 | -0.01 | -0.01 | -0.04 | 0.04 | -0.06 | -0.07 | -0.01 | 0.14 | 0.05 | -0.02 | 0.02 | -0.07 | -0.13 | -0.12 | 0.01 | 0.32 | -0.02 | -0.02 | 0.13 | -0.08 | -0.83 | 0.00 | 0.00 |
| BIO12 | 0.23 | 0.21 | 0.21 | 0.11 | 0.14 | -0.03 | 0.08 | 0.11 | -0.07 | -0.13 | 0.04 | -0.02 | 0.02 | -0.33 | -0.08 | -0.02 | 0.68 | -0.12 | 0.38 | 0.20 | -0.01 | -0.03 | -0.01 | 0.00 | 0.00 |
| BIO13 | 0.24 | 0.07 | 0.35 | -0.19 | 0.19 | -0.03 | 0.14 | 0.14 | -0.10 | -0.35 | 0.02 | -0.03 | -0.11 | 0.02 | 0.08 | 0.00 | -0.57 | 0.13 | 0.44 | 0.11 | 0.03 | 0.00 | 0.01 | 0.00 | 0.00 |
| BIO14 | 0.09 | 0.29 | -0.21 | 0.31 | 0.16 | -0.12 | 0.14 | -0.01 | -0.30 | -0.02 | 0.00 | -0.10 | -0.07 | 0.22 | -0.34 | 0.04 | -0.12 | 0.07 | -0.28 | 0.54 | -0.17 | -0.17 | 0.01 | 0.00 | 0.00 |
| BIO15 | -0.02 | -0.31 | 0.17 | -0.37 | -0.04 | -0.04 | -0.14 | -0.04 | -0.15 | -0.13 | 0.05 | -0.06 | -0.15 | 0.60 | -0.43 | 0.05 | 0.27 | -0.13 | 0.10 | -0.03 | 0.00 | 0.03 | 0.00 | 0.00 | 0.00 |

| | | | | | | | | | | | | | | | | | | | | | | | | | |
|---|---|---|---|---|---|---|---|---|---|---|---|---|---|---|---|---|---|---|---|---|---|---|---|---|---|
| BIO16 | 0.24 | 0.08 | 0.37 | -0.15 | 0.17 | -0.01 | 0.10 | 0.14 | -0.05 | -0.30 | 0.04 | -0.01 | -0.07 | -0.11 | 0.05 | -0.04 | 0.09 | -0.06 | -0.72 | -0.26 | -0.04 | -0.02 | 0.00 | 0.00 | 0.00 |
| BIO17 | 0.10 | 0.30 | -0.19 | 0.31 | 0.15 | -0.10 | 0.14 | 0.00 | -0.26 | 0.01 | -0.01 | -0.06 | -0.04 | 0.13 | -0.23 | 0.02 | -0.04 | 0.00 | 0.17 | -0.70 | 0.15 | 0.20 | 0.00 | 0.00 | 0.00 |
| Bulk density | -0.08 | -0.30 | -0.14 | 0.14 | -0.09 | -0.19 | 0.07 | -0.14 | -0.18 | -0.44 | 0.54 | -0.03 | 0.52 | -0.10 | 0.00 | -0.01 | -0.03 | 0.00 | 0.00 | -0.01 | 0.00 | 0.00 | 0.00 | 0.00 | 0.00 |
| Corse fragments | -0.05 | -0.02 | -0.39 | -0.52 | 0.14 | 0.22 | 0.23 | 0.34 | -0.37 | 0.05 | -0.24 | 0.03 | 0.33 | -0.12 | -0.01 | 0.03 | 0.06 | 0.08 | 0.00 | 0.01 | -0.01 | 0.02 | 0.00 | 0.00 | 0.00 |
| Nitrogen | -0.03 | 0.30 | 0.04 | -0.12 | 0.07 | 0.37 | -0.11 | -0.35 | -0.17 | -0.01 | 0.27 | 0.71 | -0.02 | 0.05 | -0.02 | 0.00 | 0.00 | 0.01 | 0.00 | 0.02 | 0.01 | 0.01 | 0.00 | 0.00 | 0.00 |
| pH | -0.22 | -0.18 | -0.26 | -0.06 | -0.03 | 0.10 | -0.08 | -0.17 | -0.30 | -0.26 | 0.05 | -0.15 | -0.66 | -0.42 | -0.03 | -0.05 | 0.00 | 0.01 | -0.01 | -0.01 | 0.04 | -0.02 | 0.00 | 0.00 | 0.00 |
| Sand | -0.07 | -0.28 | -0.02 | 0.13 | 0.58 | 0.04 | -0.07 | 0.00 | 0.08 | 0.05 | -0.01 | 0.07 | 0.00 | 0.01 | 0.01 | -0.01 | 0.01 | -0.01 | 0.01 | 0.00 | 0.00 | 0.00 | 0.00 | 0.74 | 0.00 |
| Silt | 0.13 | 0.19 | -0.05 | 0.08 | -0.60 | 0.05 | -0.30 | 0.35 | -0.13 | -0.22 | -0.07 | 0.09 | 0.00 | 0.03 | 0.00 | 0.01 | -0.02 | 0.02 | -0.01 | -0.01 | -0.01 | 0.00 | 0.00 | 0.52 | 0.00 |
| Clay | -0.05 | 0.25 | 0.10 | -0.32 | -0.26 | -0.13 | 0.50 | -0.44 | 0.02 | 0.18 | 0.11 | -0.24 | -0.01 | -0.06 | -0.02 | 0.01 | 0.02 | 0.00 | 0.00 | 0.00 | 0.00 | 0.01 | 0.00 | 0.42 | 0.00 |
| SOC | 0.00 | 0.29 | 0.04 | -0.07 | 0.15 | 0.44 | -0.45 | -0.22 | -0.07 | -0.05 | 0.08 | -0.60 | 0.22 | 0.07 | 0.12 | 0.01 | -0.02 | -0.02 | 0.01 | -0.01 | -0.01 | 0.00 | 0.00 | 0.00 | 0.00 |

**Table S2.** Variance explained by each Principal Component (PC) derived from 25 climatic and edaphic variables. Standard deviation of each PC, proportion of variance explained, and the cumulative proportion of variance across PCs.

| | **Standard deviation** | **Proportion of Variance** | **Cumulative Proportion** |
|---|---|---|---|
| PC1 | 3.15 | 0.40 | 0.40 |
| PC2 | 2.63 | 0.28 | 0.67 |
| PC3 | 1.41 | 0.08 | 0.75 |
| PC4 | 1.25 | 0.06 | 0.82 |
| PC5 | 1.07 | 0.05 | 0.86 |
| PC6 | 0.92 | 0.03 | 0.90 |
| PC7 | 0.74 | 0.02 | 0.92 |
| PC8 | 0.69 | 0.02 | 0.94 |
| PC9 | 0.65 | 0.02 | 0.95 |
| PC10 | 0.57 | 0.01 | 0.97 |
| PC11 | 0.51 | 0.01 | 0.98 |
| PC12 | 0.47 | 0.01 | 0.99 |
| PC13 | 0.37 | 0.01 | 0.99 |
| PC14 | 0.29 | 0.00 | 0.99 |
| PC15 | 0.25 | 0.00 | 1.00 |
| PC16 | 0.17 | 0.00 | 1.00 |
| PC17 | 0.13 | 0.00 | 1.00 |
| PC18 | 0.12 | 0.00 | 1.00 |
| PC19 | 0.08 | 0.00 | 1.00 |
| PC20 | 0.05 | 0.00 | 1.00 |
| PC21 | 0.05 | 0.00 | 1.00 |
| PC22 | 0.04 | 0.00 | 1.00 |
| PC23 | 0.02 | 0.00 | 1.00 |
| PC24 | 0.01 | 0.00 | 1.00 |
| PC25 | 0.00 | 0.00 | 1.00 |

**Table S3.** List of seed suppliers that participate in the Redário Coalition, including their municipality, state, geographic coordinates, and number of collectors per supplier.

| Supplier | Municipality | State | Latitude | Longitude | # collectors |
|---|---|---|---|---|---|
| Rede de Sementes do Xingu | Nova Xavantina | Mato Grosso | -14.6728 | -52.3528 | 647 |
| Rede de Sementes do Xingu | Água Boa, Mato Grosso | Mato Grosso | -14.0500 | -52.1589 | 647 |
| Rede de Sementes do Xingu | Canarana, Mato Grosso | Mato Grosso | -13.5500 | -52.1658 | 647 |
| Rede de Sementes do Xingu | Querência | Mato Grosso | -12.4758 | -52.3778 | 647 |
| Rede de Sementes do Xingu | Marcelândia | Mato Grosso | -11.1328 | -54.5969 | 647 |
| Rede de Sementes do Xingu | Gaúcha do Norte | Mato Grosso | -13.2419 | -53.0800 | 647 |
| Rede de Sementes do Xingu | Bom Jesus do Araguaia | Mato Grosso | -12.1778 | -51.5058 | 647 |
| Rede de Sementes do Xingu | Serra Nova Dourada | Mato Grosso | -12.0919 | -51.4019 | 647 |
| Rede de Sementes do Xingu | Canabrava do Norte | Mato Grosso | -11.0539 | -51.8308 | 647 |
| Rede de Sementes do Xingu | Porto Alegre do Norte | Mato Grosso | -10.8778 | -51.6328 | 647 |
| Rede de Sementes do Xingu | Confresa | Mato Grosso | -10.6439 | -51.5689 | 647 |
| Rede de Sementes do Xingu | São Félix do Araguaia | Mato Grosso | -11.6169 | -50.6689 | 647 |
| Rede de Sementes do Xingu | Santa Cruz do Xingu | Mato Grosso | -10.1550 | -52.3919 | 647 |
| Rede de Sementes do Xingu | São José do Xingu | Mato Grosso | -10.8039 | -52.7439 | 647 |
| Rede de Sementes do Xingu | Feliz Natal | Mato Grosso | -12.3858 | -54.9200 | 647 |
| Rede de Sementes do Xingu | Diamantino | Mato Grosso | -14.4089 | -56.4458 | 647 |
| Rede de Sementes do Xingu | Poxoréu | | -15.82660094 | -54.40210456 | 647 |
| Rede de Sementes do Xingu | Alto Boa Vista | Mato Grosso | -11.6739 | -51.3878 | 647 |
| Rede de Sementes do Portal da Amazônia | Alta Floresta | Mato Grosso | -9.8758 | -56.0858 | 200 |
| Rede de Sementes do Portal da Amazônia | Apiacás | Mato Grosso | -9.5439 | -57.4489 | 200 |
| Rede de Sementes do Portal da Amazônia | Carlinda | Mato Grosso | -9.9578 | -55.8319 | 200 |
| Rede de Sementes do Portal da Amazônia | Colíder | Mato Grosso | -10.8128 | -55.4550 | 200 |
| Rede de Sementes do Portal da Amazônia | Nova Canaã do Norte | Mato Grosso | -10.5578 | -55.9528 | 200 |
| Rede de Sementes do Portal da Amazônia | Nova Guarita | Mato Grosso | -10.3158 | -55.4100 | 200 |
| Rede de Sementes do Portal da Amazônia | Novo Mundo | Mato Grosso | -9.9500 | -55.1978 | 200 |
| Rede de Sementes do Portal da Amazônia | Peixoto de Azevedo | Mato Grosso | -10.2289 | -54.9828 | 200 |
| Rede de Sementes do Portal da Amazônia | Terra Nova do Norte | Mato Grosso | -10.5169 | -55.2308 | 200 |
| Rede de Sementes do Araguaia (RESSEMEAR) | Santana do Araguaia | Pará | -9.5050 | -50.6250 | 70 |
| Rede de Sementes do Araguaia (RESSEMEAR) | Caseara | Tocantins | -9.2778 | -49.9558 | 70 |
| Rede de Sementes do Araguaia (RESSEMEAR) | Marianópolis do Tocantins | Tocantins | -9.7958 | -49.6539 | 70 |
| Rede de Coletores do Oeste da Bahia - ARSOBA | Luís Eduardo Magalhães | Bahia | -12.0919 | -45.8050 | 81 |
| Rede de Coletores do Oeste da Bahia - ARSOBA | Barreiras | Bahia | -12.1528 | -44.9900 | 81 |
| Rede de Coletores do Oeste da Bahia - ARSOBA | São Desidério | Bahia | -12.3628 | -44.9728 | 81 |

| | | | | | |
|---|---|---|---|---|---|
| Rede de Coletores do Oeste da Bahia - ARSOBA | Angical | Bahia | -12.0069 | -44.6939 | 81 |
| Rede de Coletores do Oeste da Bahia - ARSOBA | Formosa do Rio Preto | Bahia | -11.0478 | -45.1928 | 81 |
| Rede de Sementes Sapucuá | Oriximiná | Pará | -1.7658 | -55.8658 | 60 |
| Rede de Sementes Serra da Lua (RESSEL) | Boa Vista, Roraima | Roraima | 2.8200 | -60.6719 | |
| Rede de sementes Vista Alegre | Curaçá | Bahia | -8.9919 | -39.9078 | |
| Rede de Sementes do Pajeú | Floresta, Pernambuco | Pernambuco | -8.6008 | -38.5678 | 4 |
| Catadores de Sementes do Vale do São Francisco do Sertão Baiano | Casa Nova | Bahia | -9.1619 | -40.9708 | 12 |
| Mãos que Semeiam | Assu, Rio Grande do Norte | Rio Grande do Norte | -5.5769 | -36.9089 | 9 |
| Mãos que Semeiam | Upanema | Rio Grande do Norte | -5.6419 | -37.2578 | 9 |
| Verde Novo | Mambaí | Goiás | -14.4878 | -46.1128 | 31 |
| Verde Novo | Cristalina | Goiás | -16.7689 | -47.6139 | 31 |
| Verde Novo | Arraias | Tocantins | -12.9308 | -46.9378 | 31 |
| Verde Novo | Brasília | Federal District | -15.7939 | -47.8828 | 31 |
| Verde Novo | Santa Fé de Minas | Minas Gerais | -16.6900 | -45.4139 | 31 |
| Verde Novo | Lago da Pedra | Maranhão | -4.5700 | -45.1319 | 31 |
| Aprospera | Planaltina, Federal District | Federal District | -15.6192 | -47.6525 | 17 |
| Aprospera | Formosa, Goiás | Goiás | -15.5369 | -47.3339 | 17 |
| Aprospera | Brasília | Federal District | -15.7939 | -47.8828 | 17 |
| Grupo de Coletores Geraizeiros | Montezuma, Minas Gerais | Minas Gerais | -15.1719 | -42.4969 | 56 |
| Grupo de Coletores Geraizeiros | Vargem Grande do Rio Pardo | Minas Gerais | -15.4028 | -42.3078 | 56 |
| Grupo de Coletores Geraizeiros | Rio Pardo de Minas | Minas Gerais | -15.6100 | -42.5400 | 56 |
| Grupo de Coletores Geraizeiros | Taiobeiras | Minas Gerais | -15.8078 | -42.2328 | 56 |
| Grupo de Coletores Geraizeiros | Berizal | Minas Gerais | -15.6128 | -41.7450 | 56 |
| Cooperaçú | Januária | Minas Gerais | -15.4878 | -44.3619 | 64 |
| Cooperaçú | Itacarambi | Minas Gerais | -15.1019 | -44.0919 | 64 |
| Cooperaçú | São João das Missões | Minas Gerais | -14.8839 | -44.0908 | 64 |
| Cooperaçú | Miravânia | Minas Gerais | -14.7408 | -44.4039 | 64 |
| Cooperaçú | Bonito de Minas | Minas Gerais | -15.3228 | -44.7539 | 64 |
| Sementes do Paraíso | São João do Paraíso | Minas Gerais | -15.3139 | -42.0139 | 16 |
| Sementes do Paraíso | Ninheira | Minas Gerais | -15.3208 | -41.7539 | 16 |
| Sementes do Paraíso | Rio Pardo de Minas | Minas Gerais | -15.6100 | -42.5400 | 16 |
| Rede de Cantinas da Terra do Meio | Altamira, Pará | Pará | -3.2028 | -52.2058 | 83 |
| Rede de Cantinas da Terra do Meio | Uruará | Pará | -3.7178 | -53.7369 | 83 |
| Sementes da Floresta Sul do Pará | Tucumã, Pará | Pará | -6.7519 | -51.1539 | |
| Sementes da Floresta Sul do Pará | São Félix do Xingu | Pará | -6.6450 | -51.9950 | |
| Mutum Sementes Amazônicas | Porto Velho | Rondônia | -8.7619 | -63.9039 | 50 |
| ReSeBa da Bioeconomia Amazônica (Ecoporé) | Rolim de Moura | Rondônia | -11.7254 | -61.7778 | 590 |
| ReSeBa da Bioeconomia Amazônica (Ecoporé) | Porto Velho | Rondônia | -8.7619 | -63.9039 | 590 |

| | | | | | |
|---|---|---|---|---|---|
| ReSeBa da Bioeconomia Amazônica (Ecoporé) | Alta Floresta | Mato Grosso | -9.8758 | -56.0858 | 590 |
| ReSeBa da Bioeconomia Amazônica (Ecoporé) | Alvorada d'Oeste | Rondônia | -11.3417 | -62.2861 | 590 |
| ReSeBa da Bioeconomia Amazônica (Ecoporé) | Cacoal | Rondônia | -11.4386 | -61.4472 | 590 |
| ReSeBa da Bioeconomia Amazônica (Ecoporé) | Candeias do Jamari | Rondônia | -8.8097 | -63.6956 | 590 |
| ReSeBa da Bioeconomia Amazônica (Ecoporé) | Chupinguaia | Rondônia | -12.5522 | -60.8997 | 590 |
| ReSeBa da Bioeconomia Amazônica (Ecoporé) | Costa Marques | Rondônia | -12.4450 | -64.2272 | 590 |
| ReSeBa da Bioeconomia Amazônica (Ecoporé) | Guajará-Mirim | Rondônia | -10.7828 | -65.3394 | 590 |
| ReSeBa da Bioeconomia Amazônica (Ecoporé) | Itapuã do Oeste | Rondônia | -9.2022 | -63.1800 | 590 |
| ReSeBa da Bioeconomia Amazônica (Ecoporé) | Ji-Paraná | Rondônia | -10.8853 | -61.9517 | 590 |
| ReSeBa da Bioeconomia Amazônica (Ecoporé) | Nova Brasilândia d'Oeste | Rondônia | -11.7197 | -62.3158 | 590 |
| ReSeBa da Bioeconomia Amazônica (Ecoporé) | Presidente Médici, Rondônia | Rondônia | -11.1758 | -61.9008 | 590 |
| ReSeBa da Bioeconomia Amazônica (Ecoporé) | Rondolândia | Mato Grosso | -10.5000 | -61.0000 | 590 |
| ReSeBa da Bioeconomia Amazônica (Ecoporé) | Santa Luzia d'Oeste | Rondônia | -11.9081 | -61.7789 | 590 |
| ReSeBa da Bioeconomia Amazônica (Ecoporé) | São Felipe d'Oeste | Rondônia | -11.9025 | -61.5022 | 590 |
| ReSeBa da Bioeconomia Amazônica (Ecoporé) | São Francisco do Guaporé | Rondônia | -12.0522 | -63.5675 | 590 |
| ReSeBa da Bioeconomia Amazônica (Ecoporé) | São Miguel do Guaporé | Rondônia | -11.6936 | -62.7114 | 590 |
| ReSeBa da Bioeconomia Amazônica (Ecoporé) | Seringueiras | Rondônia | -11.7981 | -63.0311 | 590 |
| ReSeBa da Bioeconomia Amazônica (Ecoporé) | Urupá | Rondônia | -11.1406 | -62.3608 | 590 |
| ReSeBa da Bioeconomia Amazônica (Ecoporé) | Vilhena | Rondônia | -12.7406 | -60.1458 | 590 |
| Coopajé | Porto Seguro | Bahia | -16.4500 | -39.0650 | 20 |
| Coopajé | Itamaraju | Bahia | -17.0389 | -39.5308 | 20 |
| Coopajé | Camacan | Bahia | -15.4189 | -39.4958 | 20 |
| Coopajé | Pau Brasil | Bahia | -15.4639 | -39.6508 | 20 |
| Coopajé | Itabela | Bahia | -16.5750 | -39.5528 | 20 |
| Rede Primaflora | Prado, Bahia | Bahia | -17.3408 | -39.2208 | 126 |
| Rede Primaflora | Mucuri | Bahia | -18.0858 | -39.5508 | 126 |
| Rede Primaflora | Medeiros Neto | Bahia | -17.3739 | -40.2208 | 126 |
| Rede Primaflora | Itamaraju | Bahia | -17.0389 | -39.5308 | 126 |
| Rede Primaflora | Nova Viçosa | Bahia | -17.8919 | -39.3719 | 126 |
| Rede Primaflora | Itamaraju | Bahia | -17.0389 | -39.5308 | 126 |
| Rede Primaflora | Caravelas | Bahia | -17.7319 | -39.2658 | 126 |
| Rede Primaflora | Porto Seguro | Bahia | -16.4500 | -39.0650 | 126 |
| Rede Primaflora | Belmonte, Bahia | Bahia | -15.8628 | -38.8828 | 126 |
| Rede Primaflora | Ilhéus | Bahia | -14.7889 | -39.0489 | 126 |
| Rede Primaflora | Serra Grande, Paraíba | Paraíba | -7.2150 | -38.3700 | 126 |
| Rede Primaflora | Jacinto, Minas Gerais | Minas Gerais | -16.1439 | -40.2928 | 126 |
| Rede Primaflora | Itaobim | Minas Gerais | -16.5619 | -41.5028 | 126 |
| Rede Primaflora | Pocrane | Minas Gerais | -19.6200 | -41.6369 | 126 |

| | | | | | |
|---|---|---|---|---|---|
| Rede Primaflora | Nanuque | Minas Gerais | -17.8389 | -40.3539 | 126 |
| Rede Primaflora | Montanha, Espírito Santo | Espírito Santo | -18.1269 | -40.3628 | 126 |
| Rede Primaflora | São Gabriel da Palha | Espírito Santo | -19.0169 | -40.5358 | 126 |
| Rede Primaflora | Guarapari | Espírito Santo | -20.6578 | -40.5108 | 126 |
| Rede Primaflora | Itamaraju | Bahia | -17.0389 | -39.5308 | 126 |
| Rede Primaflora | Itamaraju | Bahia | -17.0389 | -39.5308 | 126 |
| Rede Primaflora | Itamaraju | Bahia | -17.0389 | -39.5308 | 126 |
| Arboretum | Teixeira de Freitas | Bahia | -17.5350 | -39.7419 | 23 |
| Arboretum | Caravelas | Bahia | -17.7319 | -39.2658 | 23 |
| Arboretum | Itamaraju | Bahia | -17.0389 | -39.5308 | 23 |
| Arboretum | Porto Seguro | Bahia | -16.4500 | -39.0650 | 23 |
| Arboretum | Mucuri | Bahia | -18.0858 | -39.5508 | 23 |
| Rede Tupyguá | Aracruz, Espírito Santo | Espírito Santo | -19.8200 | -40.2728 | 52 |
| Rede Tupyguá | Linhares | Espírito Santo | -19.3987 | -40.0651 | 52 |
| Refloresta | Boituva | São Paulo | -23.2833 | -47.6722 | 3 |
| Refloresta | Capivari | São Paulo | -22.9950 | -47.5078 | 3 |
| Refloresta | Cerquilho | São Paulo | -23.1650 | -47.7436 | 3 |
| Refloresta | Porto Feliz | São Paulo | -23.2150 | -47.5239 | 3 |
| Refloresta | Rafard | São Paulo | -23.0117 | -47.5269 | 3 |
| Refloresta | Sorocaba | São Paulo | -23.5019 | -47.4578 | 3 |
| Refloresta | Tietê, São Paulo | São Paulo | -23.1019 | -47.7150 | 3 |
| Refloresta | Itapirapuã Paulista | São Paulo | -24.5739 | -49.2969 | 3 |
| Refloresta | Piedade, São Paulo | São Paulo | -23.7119 | -47.4278 | 3 |
| Refloresta | Pilar do Sul | São Paulo | -23.8128 | -47.7158 | 3 |
| Refloresta | Ribeirão Grande | São Paulo | -24.0989 | -48.3650 | 3 |
| Refloresta | São Miguel Arcanjo | São Paulo | -23.8778 | -47.9969 | 3 |
| Refloresta | Itu, São Paulo | São Paulo | -23.2642 | -47.2992 | 3 |
| Refloresta | Buri, São Paulo | São Paulo | -23.7975 | -48.5928 | 3 |
| Refloresta | Capão Bonito | São Paulo | -24.0058 | -48.3494 | 3 |
| Refloresta | Guapiara | São Paulo | -24.1850 | -48.5328 | 3 |
| Refloresta | Itapetininga | São Paulo | -23.5917 | -48.0531 | 3 |
| Refloresta | Itapeva, São Paulo | São Paulo | -23.9819 | -48.8758 | 3 |
| Rede de Sementes do Ribeira | Eldorado, São Paulo | São Paulo | -24.5200 | -48.1081 | 64 |
| Rede de Sementes do Ribeira | Iporanga | São Paulo | -24.5856 | -48.5931 | 64 |
| Rede de Coletores do Vale do Paraíba | Lagoinha | São Paulo | -23.0906 | -45.1903 | 75 |
| Rede de Coletores do Vale do Paraíba | Cunha, São Paulo | São Paulo | -23.0744 | -44.9597 | 75 |
| Rede de Coletores do Vale do Paraíba | São Luiz do Paraitinga | São Paulo | -23.2219 | -45.3100 | 75 |
| Rede de Coletores do Vale do Paraíba | Taubaté | São Paulo | -23.0333 | -45.5500 | 75 |

| | | | | | |
|---|---|---|---|---|---|
| Rede de Coletores do Vale do Paraíba | Pindamonhangaba | São Paulo | -22.9239 | -45.4617 | 75 |
| Rede de Coletores do Vale do Paraíba | Tremembé | São Paulo | -22.9578 | -45.5489 | 75 |
| Rede de Coletores do Vale do Paraíba | São José dos Campos | São Paulo | -23.1789 | -45.8869 | 75 |
| Rede de Coletores do Vale do Paraíba | Caçapava | São Paulo | -23.1008 | -45.7069 | 75 |
| Rede de Coletores do Vale do Paraíba | Cruzeiro, São Paulo | São Paulo | -22.5772 | -44.9583 | 75 |
| Rede de Coletores do Vale do Paraíba | Jambeiro | São Paulo | -23.2536 | -45.6878 | 75 |
| Flora Tietê | Penápolis | São Paulo | -21.4200 | -50.0778 | 2 |
| Flora Tietê | Araçatuba | São Paulo | -21.2089 | -50.4328 | 2 |
| Flora Tietê | Sales, São Paulo | São Paulo | -21.3408 | -49.4850 | 2 |
| Flora Tietê | São José do Rio Preto | São Paulo | -20.8200 | -49.3789 | 2 |
| Flora Tietê | Promissão | São Paulo | -21.5369 | -49.8583 | 2 |
| Flora Tietê | José Bonifácio, São Paulo | São Paulo | -21.0528 | -49.6878 | 2 |
| Zé da Lena Sementes | Laranjal Paulista | São Paulo | -23.0515 | -47.8370 | 6 |
| Zé da Lena Sementes | Santa Bárbara d'Oeste | São Paulo | -22.7539 | -47.4139 | 6 |
| Zé da Lena Sementes | Botucatu | São Paulo | -22.8858 | -48.4450 | 6 |
| Zé da Lena Sementes | Conchas | São Paulo | -23.0134 | -48.0078 | 6 |
| Zé da Lena Sementes | Piracicaba | São Paulo | -22.7252 | -47.6493 | 6 |
| Zé da Lena Sementes | Anhembi | São Paulo | -22.7897 | -48.1278 | 6 |
| Zé da Lena Sementes | Itu, São Paulo | São Paulo | -23.2642 | -47.2992 | 6 |
| Zé da Lena Sementes | Penápolis | São Paulo | -21.4200 | -50.0778 | 6 |
| Zé da Lena Sementes | Sumaré | São Paulo | -22.8219 | -47.2669 | 6 |
| Associação Cerrado de Pé | Alto Paraíso de Goiás | Goiás | -14.1328 | -47.5100 | 200 |
| Associação Cerrado de Pé | Cavalcante | Goiás | -13.7978 | -47.4578 | 200 |
| Associação Cerrado de Pé | Teresina de Goiás | Goiás | -13.7758 | -47.2650 | 200 |
| Associação Cerrado de Pé | Colinas do Sul | Goiás | -14.1508 | -48.0778 | 200 |
| Monjolinho | São Carlos | São Paulo | -22.0178 | -47.8908 | 1 |
| Rede de Sementes +Floresta | Abelardo Luz | Santa Catarina | -26.5650 | -52.3278 | 12 |
| Rede de Sementes e Mudas da Bacia do São Francisco | Petrolina | Pernambuco | -9.3928 | -40.5078 | |
| Rede de Sementes e Mudas da Bacia do São Francisco | Juazeiro | Bahia | -9.4139 | -40.5028 | |

**Table S4.** Characteristics of the 48 Seed Transfer Zones (STZs) in Brazil. For each STZ, the total area and its proportion of Brazil's territory, number of seed suppliers and seed collection sites, number of seed collectors and their density per 100,000 km$^2$, native vegetation relativized by area, legal restoration deficit relativized by area, degraded pasture coverage relativized by area, projected STZ area for 2041–2060 and 2081–2100 under two future climate scenarios (SSP1 and SSP5) based on the MPI-ESM1.2-HR climate model, and the relative change in area (ΔArea) for each timeframe and under each scenario (Future - Present).

| STZ | Area (km²) | National coverage (%) | Seed suppliers (no.) | Seed collection sites (no.) | Number of collectors (no.) | Collector density (no. per 100,000 km²) | Native vegetation (%) | Legal deficit (%) | Degraded pasture (%) | Area 2060 SSP1 (km²) | Area 2060 SSP5 (km²) | Area 2100 SSP1 (km²) | Area 2100 SSP5 (km²) | ΔArea 2060 SSP1 (%) | ΔArea 2060 SSP5 (%) | ΔArea 2100 SSP1 (%) | ΔArea 2100 SSP5 (%) |
|---|---|---|---|---|---|---|---|---|---|---|---|---|---|---|---|---|---|
| 1 | 73288.75 | 0.86 | 0.00 | 0.00 | 0.00 | 0.00 | 78.17 | 0.62 | 0.45 | 79356.74 | 78084.18 | 71789.55 | 95957.71 | 8.28 | 6.54 | -2.05 | 30.93 |
| 2 | 58151.60 | 0.68 | 0.00 | 0.00 | 0.00 | 0.00 | 93.81 | 0.18 | 0.04 | 51550.73 | 23043.17 | 49750.55 | 15404.45 | -11.35 | -60.37 | -14.45 | -73.51 |
| 3 | 369476.64 | 4.32 | 0.00 | 0.00 | 0.00 | 0.00 | 90.55 | 0.30 | 0.24 | 315410.45 | 250505.57 | 324187.84 | 98538.07 | -14.63 | -32.20 | -12.26 | -73.33 |
| 4 | 168388.10 | 1.97 | 1.00 | 1.00 | 1.00 | 0.53 | 89.37 | 0.35 | 0.51 | 244635.21 | 333476.05 | 263227.89 | 222840.53 | 45.28 | 98.04 | 56.32 | 32.34 |
| 5 | 197533.55 | 2.31 | 0.00 | 0.00 | 0.00 | 0.00 | 90.46 | 0.18 | 0.18 | 259944.78 | 312576.06 | 271536.77 | 128171.89 | 31.60 | 58.24 | 37.46 | -35.11 |
| 6 | 237804.82 | 2.78 | 1.00 | 1.00 | 60.00 | 22.46 | 93.32 | 0.23 | 0.11 | 202420.59 | 122817.17 | 205014.44 | 28038.11 | -14.88 | -48.35 | -13.79 | -88.21 |
| 7 | 210630.82 | 2.46 | 1.00 | 2.00 | 83.00 | 33.33 | 69.49 | 2.29 | 1.53 | 152139.73 | 76553.67 | 121367.59 | 15271.38 | -27.77 | -63.66 | -42.38 | -92.75 |
| 8 | 109897.03 | 1.28 | 0.00 | 0.00 | 0.00 | 0.00 | 61.11 | 1.48 | 2.05 | 85676.80 | 72191.70 | 88856.16 | 48260.90 | -22.04 | -34.31 | -19.15 | -56.09 |
| 9 | 122481.99 | 1.43 | 0.00 | 0.00 | 0.00 | 0.00 | 98.39 | 0.01 | 0.01 | 125577.82 | 100431.05 | 127543.34 | 7540.59 | 2.53 | -18.00 | 4.13 | -93.84 |
| 10 | 159920.16 | 1.87 | 0.00 | 0.00 | 0.00 | 0.00 | 96.88 | 0.01 | 0.01 | 137956.16 | 137039.03 | 129560.60 | 169810.56 | -13.73 | -14.31 | -18.98 | 6.18 |
| 11 | 207701.13 | 2.43 | 0.00 | 0.00 | 0.00 | 0.00 | 91.29 | 0.05 | 0.20 | 240102.33 | 355906.18 | 248078.08 | 738654.43 | 15.60 | 71.35 | 19.44 | 255.63 |
| 12 | 221683.10 | 2.59 | 0.00 | 0.00 | 0.00 | 0.00 | 95.20 | 0.01 | 0.06 | 237404.28 | 174647.18 | 237250.30 | 72039.72 | 7.09 | -21.22 | 7.02 | -67.50 |
| 13 | 104568.18 | 1.22 | 1.00 | 1.00 | 5.17 | 3.38 | 52.98 | 2.89 | 3.07 | 195933.10 | 369043.41 | 209005.42 | 1309185.07 | 87.37 | 252.92 | 99.87 | 1151.99 |
| 14 | 182057.00 | 2.13 | 0.00 | 0.00 | 0.00 | 0.00 | 76.42 | 2.96 | 0.97 | 74342.98 | 13460.84 | 70957.21 | 11847.83 | -59.16 | -92.61 | -61.02 | -93.49 |
| 15 | 241605.14 | 2.82 | 2.00 | 3.00 | 15.00 | 3.97 | 62.07 | 0.52 | 5.51 | 245862.81 | 280054.15 | 244338.89 | 412778.45 | 1.76 | 15.91 | 1.13 | 70.85 |
| 16 | 181871.84 | 2.12 | 0.00 | 0.00 | 0.00 | 0.00 | 68.50 | 1.59 | 1.87 | 214527.14 | 142859.59 | 214181.14 | 68061.09 | 17.96 | -21.45 | 17.76 | -62.58 |
| 17 | 229354.22 | 2.68 | 0.00 | 0.00 | 0.00 | 0.00 | 97.74 | 0.08 | 0.02 | 168976.54 | 107115.97 | 172174.18 | 767.79 | -26.33 | -53.30 | -24.93 | -99.67 |
| 18 | 142569.46 | 1.67 | 1.00 | 2.00 | 12.00 | 4.79 | 48.47 | 1.16 | 3.71 | 57168.16 | 38602.39 | 67871.93 | 10000.58 | -59.90 | -72.92 | -52.39 | -92.99 |
| 19 | 171273.11 | 2.00 | 0.00 | 0.00 | 0.00 | 0.00 | 80.11 | 0.79 | 0.93 | 201630.86 | 234826.75 | 202969.65 | 276529.46 | 17.72 | 37.11 | 18.51 | 61.46 |
| 20 | 224795.76 | 2.63 | 1.00 | 3.00 | 70.00 | 24.17 | 46.71 | 5.37 | 7.33 | 356939.40 | 459170.39 | 398891.70 | 228256.59 | 58.78 | 104.26 | 77.45 | 1.54 |
| 21 | 131071.04 | 1.53 | 2.00 | 2.00 | 36.44 | 24.60 | 83.38 | 2.50 | 2.57 | 110134.96 | 110032.90 | 102940.81 | 73542.54 | -15.97 | -16.05 | -21.46 | -43.89 |

| | | | | | | | | | | | | | | | | | |
|---|---|---|---|---|---|---|---|---|---|---|---|---|---|---|---|---|---|
| 22 | 215508.18 | 2.52 | 2.00 | 3.00 | 106.19 | 45.12 | 94.23 | 0.54 | 0.50 | 338706.69 | 394521.43 | 323480.05 | 526337.72 | 57.17 | 83.07 | 50.10 | 144.23 |
| 23 | 332199.08 | 3.88 | 3.00 | 3.00 | 58.67 | 16.25 | 83.80 | 1.43 | 1.51 | 354622.19 | 297958.53 | 331649.67 | 203254.21 | 6.75 | -10.31 | -0.17 | -38.82 |
| 24 | 147140.91 | 1.72 | 4.00 | 5.00 | 18.00 | 9.43 | 59.85 | 0.31 | 6.56 | 174657.83 | 239934.88 | 157109.79 | 321588.19 | 18.70 | 63.06 | 6.78 | 118.56 |
| 25 | 235745.57 | 2.75 | 3.00 | 17.00 | 87.80 | 24.54 | 37.41 | 1.20 | 11.21 | 258542.60 | 241954.11 | 244679.66 | 131588.00 | 9.67 | 2.63 | 3.79 | -44.18 |
| 26 | 163390.85 | 1.91 | 2.00 | 3.00 | 45.20 | 22.43 | 73.11 | 0.97 | 2.87 | 120187.08 | 135968.06 | 125815.56 | 74902.26 | -26.44 | -16.78 | -23.00 | -54.16 |
| 27 | 219456.93 | 2.56 | 3.00 | 9.00 | 110.13 | 37.36 | 53.25 | 1.11 | 11.05 | 243750.64 | 259412.60 | 214996.28 | 209386.88 | 11.07 | 18.21 | -2.03 | -4.59 |
| 28 | 129441.63 | 1.51 | 1.00 | 1.00 | 28.10 | 18.51 | 70.90 | 3.95 | 4.88 | 83426.01 | 9107.55 | 88601.71 | | -35.55 | -92.96 | -31.55 | |
| 29 | 292179.64 | 3.41 | 3.00 | 24.00 | 643.00 | 181.51 | 58.45 | 3.56 | 5.26 | 243825.88 | 166755.60 | 253598.48 | 7148.57 | -16.55 | -42.93 | -13.20 | -97.55 |
| 30 | 189637.10 | 2.22 | 0.00 | 0.00 | 0.00 | 0.00 | 84.48 | 1.35 | 0.40 | 145751.44 | 108807.54 | 134448.62 | 44358.25 | -23.14 | -42.62 | -29.10 | -76.61 |
| 31 | 218359.20 | 2.55 | 1.00 | 8.00 | 287.56 | 102.45 | 47.03 | 3.07 | 15.83 | 367014.51 | 654868.24 | 346163.42 | 1125638.64 | 68.08 | 199.90 | 58.53 | 415.50 |
| 32 | 117044.14 | 1.37 | 1.00 | 4.00 | 112.38 | 84.69 | 73.91 | 1.55 | 2.47 | 132150.20 | 170049.20 | 134209.16 | 372081.05 | 12.91 | 45.29 | 14.67 | 217.90 |
| 33 | 194642.40 | 2.27 | 0.00 | 0.00 | 0.00 | 0.00 | 30.02 | 4.16 | 5.20 | 186735.08 | 126079.43 | 211453.93 | 117720.61 | -4.06 | -35.23 | 8.64 | -39.52 |
| 34 | 302853.63 | 3.54 | 5.00 | 10.00 | 193.87 | 45.47 | 44.03 | 1.91 | 8.24 | 246555.12 | 222602.82 | 269698.99 | 130587.26 | -18.59 | -26.50 | -10.95 | -56.88 |
| 35 | 243809.50 | 2.85 | 1.00 | 6.00 | 215.67 | 75.04 | 53.09 | 5.46 | 6.00 | 198206.65 | 120732.84 | 184826.23 | 14125.71 | -18.70 | -50.48 | -24.19 | -94.21 |
| 36 | 210137.71 | 2.45 | 5.00 | 19.00 | 170.67 | 54.40 | 43.36 | 1.94 | 12.55 | 174800.99 | 171255.27 | 189867.95 | 78100.52 | -16.82 | -18.50 | -9.65 | -62.83 |
| 37 | 103937.19 | 1.21 | 0.00 | 0.00 | 0.00 | 0.00 | 71.89 | 1.31 | 5.74 | 159352.88 | 218119.60 | 141653.32 | 299511.39 | 53.32 | 109.86 | 36.29 | 188.17 |
| 38 | 99691.87 | 1.16 | 3.00 | 13.00 | 95.17 | 57.55 | 53.71 | 0.68 | 1.05 | 90267.71 | 78206.89 | 90755.85 | 42797.23 | -9.45 | -21.55 | -8.96 | -57.07 |
| 39 | 239547.16 | 2.80 | 3.00 | 4.00 | 70.17 | 17.39 | 31.97 | 1.54 | 7.41 | 172836.78 | 115883.04 | 185517.02 | 28691.69 | -27.85 | -51.62 | -22.56 | -88.02 |
| 40 | 92313.47 | 1.08 | 4.00 | 24.00 | 38.17 | 24.06 | 17.03 | 3.03 | 2.87 | 92214.81 | 65620.17 | 88195.37 | 14567.88 | -0.11 | -28.92 | -4.46 | -84.22 |
| 41 | 166952.22 | 1.95 | 1.00 | 1.00 | 0.33 | 0.15 | 24.07 | 4.15 | 17.89 | 132169.94 | 34518.76 | 139072.93 | 772.51 | -20.83 | -79.32 | -16.70 | -99.54 |
| 42 | 121936.21 | 1.42 | 0.00 | 0.00 | 0.00 | 0.00 | 57.41 | 2.62 | 16.61 | 196808.88 | 284193.19 | 167710.99 | 250356.33 | 61.40 | 133.07 | 37.54 | 105.32 |
| 43 | 222970.74 | 2.60 | 2.00 | 6.00 | 2.33 | 0.73 | 22.89 | 3.99 | 7.54 | 175452.66 | 146976.60 | 185545.98 | 107972.72 | -21.31 | -34.08 | -16.78 | -51.58 |
| 44 | 35080.55 | 0.41 | 0.00 | 0.00 | 0.00 | 0.00 | 12.83 | 4.96 | 9.71 | 26716.02 | 22296.27 | 30445.61 | 9102.60 | -23.84 | -36.44 | -13.21 | -74.05 |
| 45 | 169906.16 | 1.98 | 0.00 | 0.00 | 0.00 | 0.00 | 25.95 | 1.49 | 0.17 | 183888.84 | 195481.10 | 188757.34 | 230433.77 | 8.23 | 15.05 | 11.10 | 35.62 |
| 46 | 196737.69 | 2.30 | 1.00 | 1.00 | 12.00 | 3.69 | 44.69 | 0.83 | 0.34 | 137367.22 | 117713.81 | 145780.13 | 55101.38 | -30.18 | -40.17 | -25.90 | -71.99 |
| 47 | 70407.08 | 0.82 | 0.00 | 0.00 | 0.00 | 0.00 | 48.40 | 1.05 | 0.12 | 50765.96 | 48028.34 | 55922.54 | 23316.41 | -27.90 | -31.78 | -20.57 | -66.88 |
| 48 | 83011.52 | 0.97 | 0.00 | 0.00 | 0.00 | 0.00 | 48.68 | 1.30 | 0.01 | 115695.60 | 120678.54 | 108711.19 | 109222.30 | 39.37 | 45.38 | 30.96 | 31.57 |

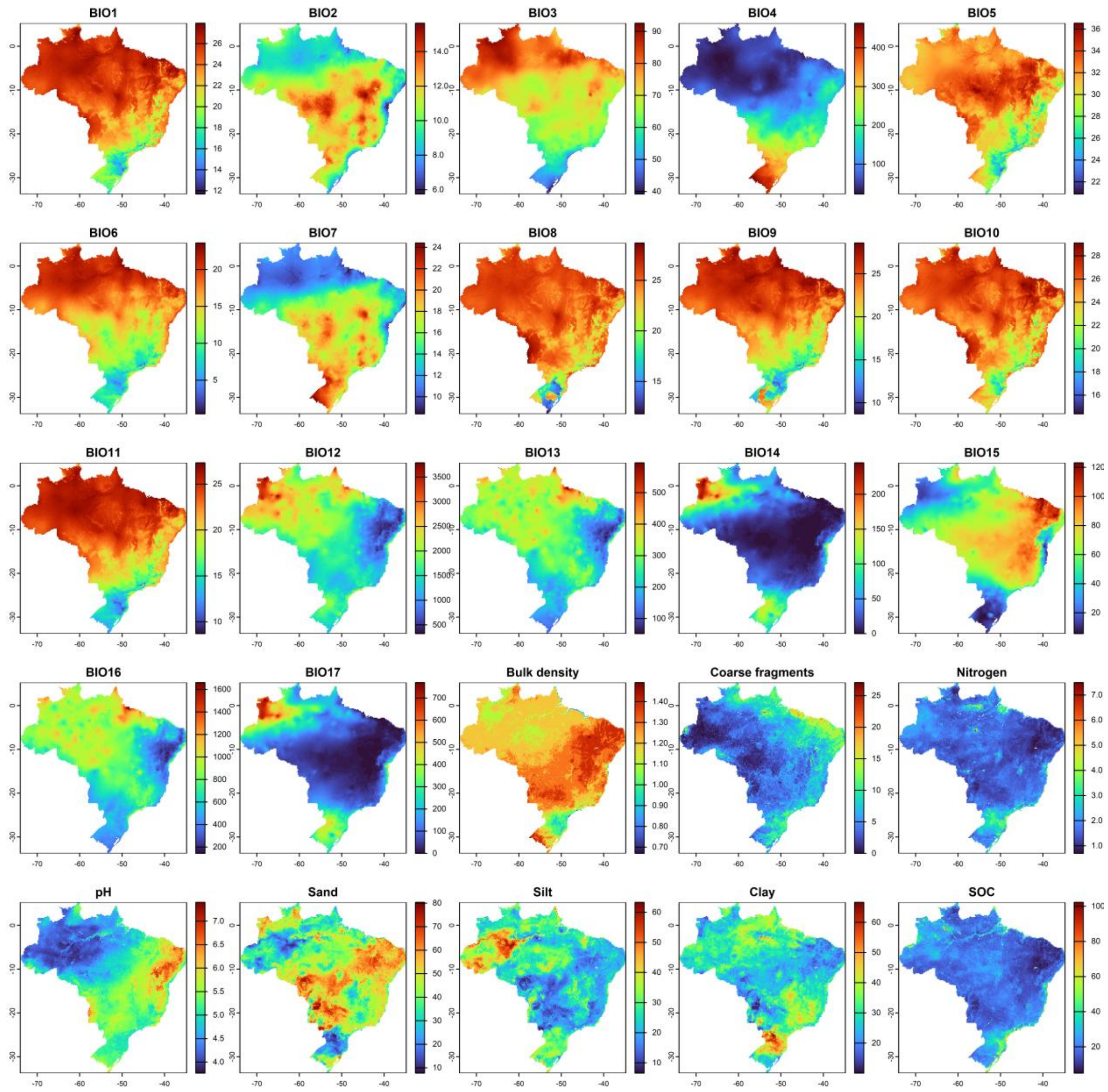


**Figure S1.** Environmental variables across Brazil. Climatic variables include BIO1, annual mean temperature (°C); BIO2, mean diurnal temperature range (°C); BIO3, isothermality (%); BIO4, temperature seasonality (standard deviation ×100, °C); BIO5, maximum temperature of the warmest month (°C); BIO6, minimum temperature of the coldest month (°C); BIO7, temperature annual range (°C); BIO8, mean temperature of the wettest quarter (°C); BIO9, mean temperature of the driest quarter (°C); BIO10, mean temperature of the warmest quarter (°C); BIO11, mean temperature of the coldest quarter (°C); BIO12, annual precipitation (mm); BIO13, precipitation of the wettest month (mm); BIO14, precipitation of the driest month (mm); BIO15, precipitation seasonality (coefficient of variation, %); BIO16, precipitation of the wettest quarter (mm); and BIO17, precipitation of the driest quarter (mm). Edaphic variables include bulk density (kg/dm$^3$), coarse fragments (%), nitrogen content (g/kg), pH (unitless), sand, silt, and clay (%), and soil organic carbon (SOC; g/kg).

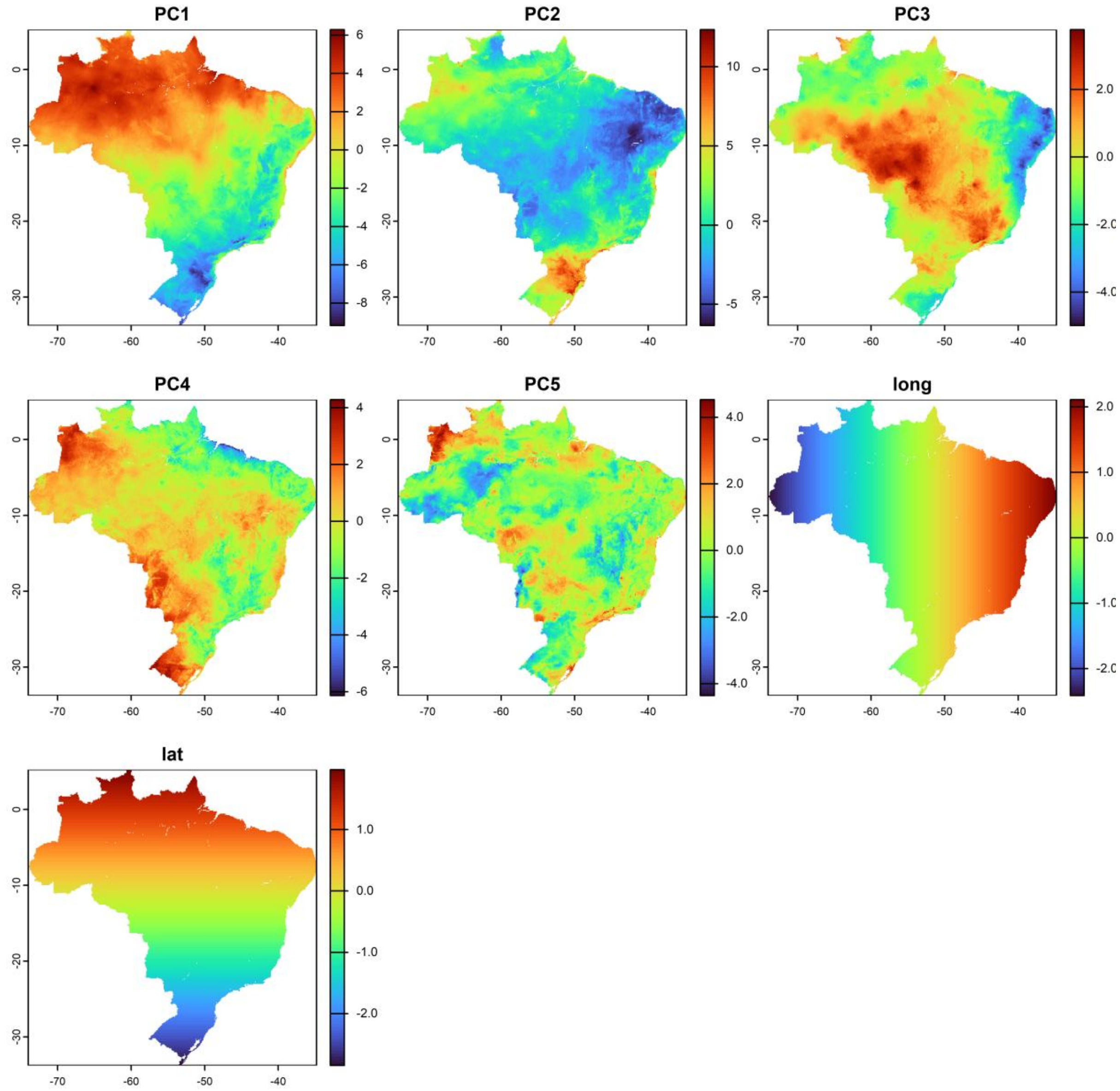


**Figure S2.** Principal Component Analysis (PCA) dimensions with eigenvalues greater than one, and longitude (long) and latitude (lat).

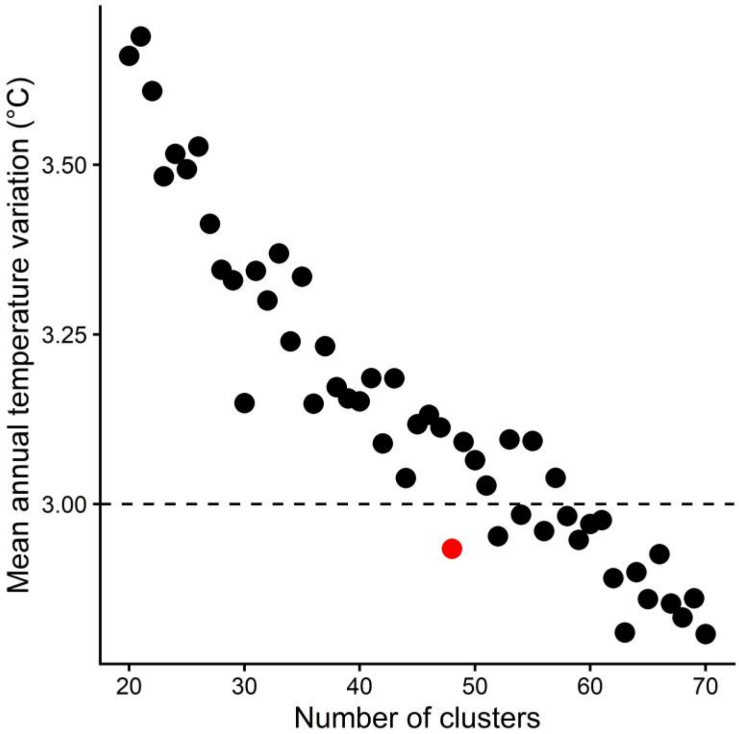


**Figure S3.** Relationship between the mean annual temperature (BIO1) range calculated within each cluster and averaged across STZ runs, plotted against the number of clusters (k). Each point represents one STZ run with k values ranging from 20 to 70. The red point indicates the STZ version with the lowest k for which the mean BIO1 range across clusters is below 3 °C.

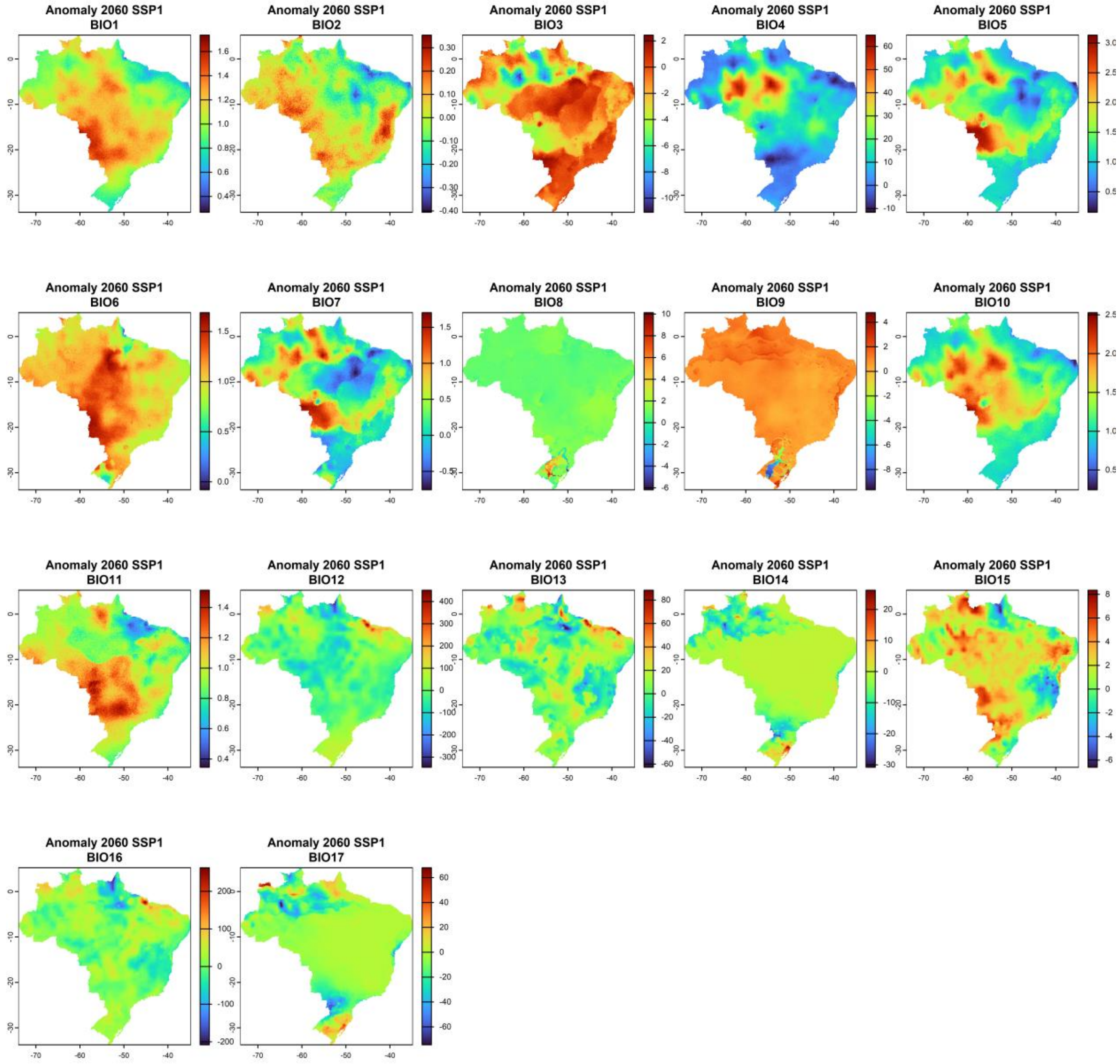


**Figure S4.** Climate anomalies under the Socioeconomic Shared Pathway 1 (SSP1), calculated as the difference between 2041–2060 and 1970–2000 averages based on the MPI-ESM1.2-HR climate model. BIO1 corresponds to annual mean temperature (°C), BIO2 to mean diurnal temperature range (°C), BIO3 to isothermality (%), BIO4 to temperature seasonality (standard deviation ×100, °C), BIO5 to maximum temperature of the warmest month (°C), BIO6 to minimum temperature of the coldest month (°C), BIO7 to temperature annual range (°C), BIO8 to mean temperature of the wettest quarter (°C), BIO9 to mean temperature of the driest quarter (°C), BIO10 to mean temperature of the warmest quarter (°C), BIO11 to mean temperature of the coldest quarter (°C), BIO12 to annual precipitation (mm), BIO13 to precipitation of the wettest month (mm), BIO14 to precipitation of the driest month (mm), BIO15 to precipitation seasonality (coefficient of variation, %), BIO16 to precipitation of the wettest quarter (mm), and BIO17 to precipitation of the driest quarter (mm).

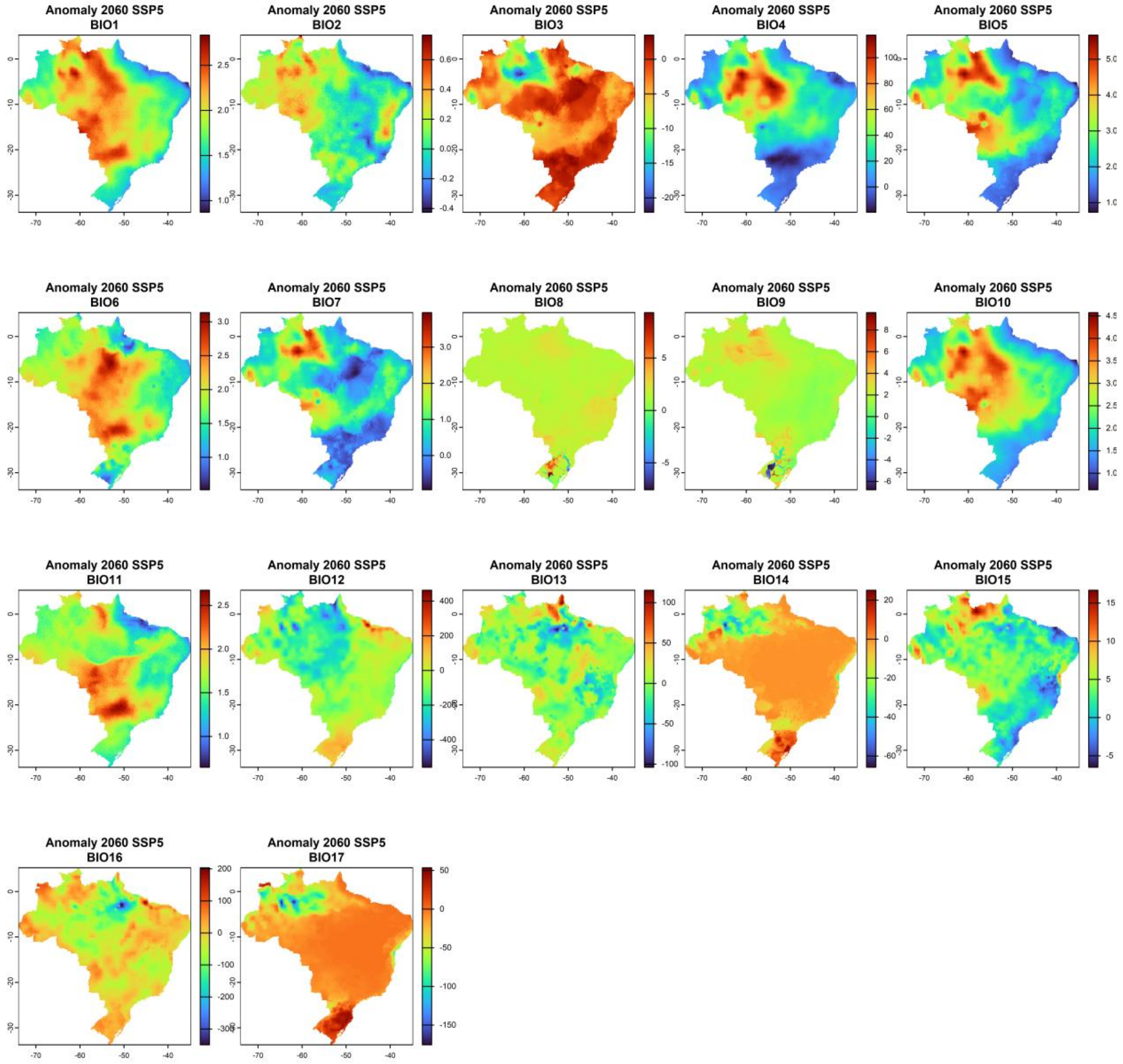


**Figure S5.** Climate anomalies under the Socioeconomic Shared Pathway 5 (SSP5), calculated as the difference between 2041–2060 and 1970–2000 averages based on the MPI-ESM1.2-HR climate model. BIO1 corresponds to annual mean temperature (°C), BIO2 to mean diurnal temperature range (°C), BIO3 to isothermality (%), BIO4 to temperature seasonality (standard deviation ×100, °C), BIO5 to maximum temperature of the warmest month (°C), BIO6 to minimum temperature of the coldest month (°C), BIO7 to temperature annual range (°C), BIO8 to mean temperature of the wettest quarter (°C), BIO9 to mean temperature of the driest quarter (°C), BIO10 to mean temperature of the warmest quarter (°C), BIO11 to mean temperature of the coldest quarter (°C), BIO12 to annual precipitation (mm), BIO13 to precipitation of the wettest month (mm), BIO14 to precipitation of the driest month (mm), BIO15 to precipitation seasonality (coefficient of variation, %), BIO16 to precipitation of the wettest quarter (mm), and BIO17 to precipitation of the driest quarter (mm).

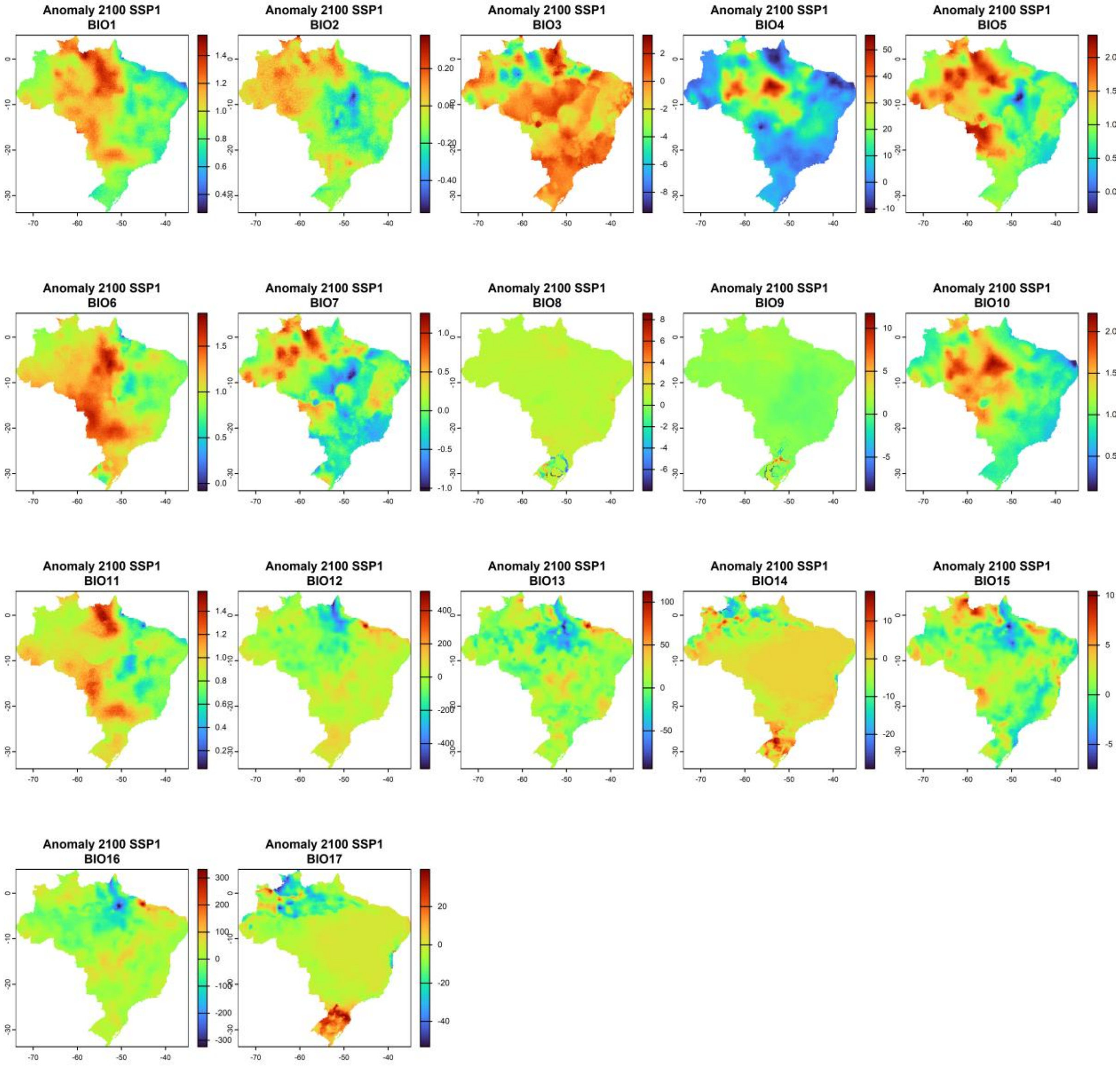


**Figure S6.** Climate anomalies under the Socioeconomic Shared Pathway 1 (SSP1), calculated as the difference between 2081–2100 and 1970–2000 averages based on the MPI-ESM1.2-HR climate model. BIO1 corresponds to annual mean temperature (°C), BIO2 to mean diurnal temperature range (°C), BIO3 to isothermality (%), BIO4 to temperature seasonality (standard deviation ×100, °C), BIO5 to maximum temperature of the warmest month (°C), BIO6 to minimum temperature of the coldest month (°C), BIO7 to temperature annual range (°C), BIO8 to mean temperature of the wettest quarter (°C), BIO9 to mean temperature of the driest quarter (°C), BIO10 to mean temperature of the warmest quarter (°C), BIO11 to mean temperature of the coldest quarter (°C), BIO12 to annual precipitation (mm), BIO13 to precipitation of the wettest month (mm), BIO14 to precipitation of the driest month (mm), BIO15 to precipitation seasonality (coefficient of variation, %), BIO16 to precipitation of the wettest quarter (mm), and BIO17 to precipitation of the driest quarter (mm).

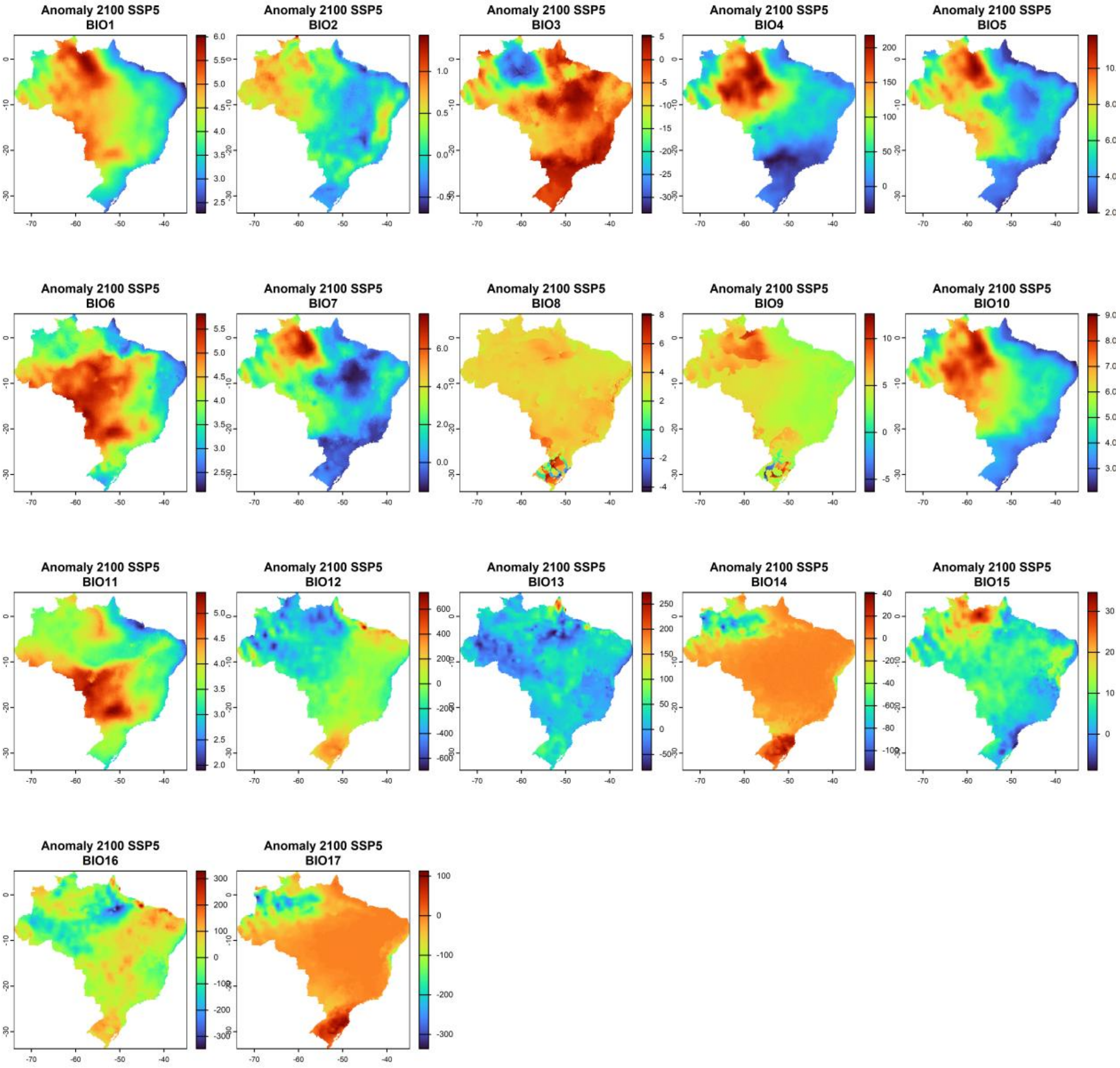


**Figure S7.** Climate anomalies under the Socioeconomic Shared Pathway 5 (SSP5), calculated as the difference between 2081–2100 and 1970–2000 averages based on the MPI-ESM1.2-HR climate model. BIO1 corresponds to annual mean temperature (°C), BIO2 to mean diurnal temperature range (°C), BIO3 to isothermality (%), BIO4 to temperature seasonality (standard deviation ×100, °C), BIO5 to maximum temperature of the warmest month (°C), BIO6 to minimum temperature of the coldest month (°C), BIO7 to temperature annual range (°C), BIO8 to mean temperature of the wettest quarter (°C), BIO9 to mean temperature of the driest quarter (°C), BIO10 to mean temperature of the warmest quarter (°C), BIO11 to mean temperature of the coldest quarter (°C), BIO12 to annual precipitation (mm), BIO13 to precipitation of the wettest month (mm), BIO14 to precipitation of the driest month (mm), BIO15 to precipitation seasonality (coefficient of variation, %), BIO16 to precipitation of the wettest quarter (mm), and BIO17 to precipitation of the driest quarter (mm).

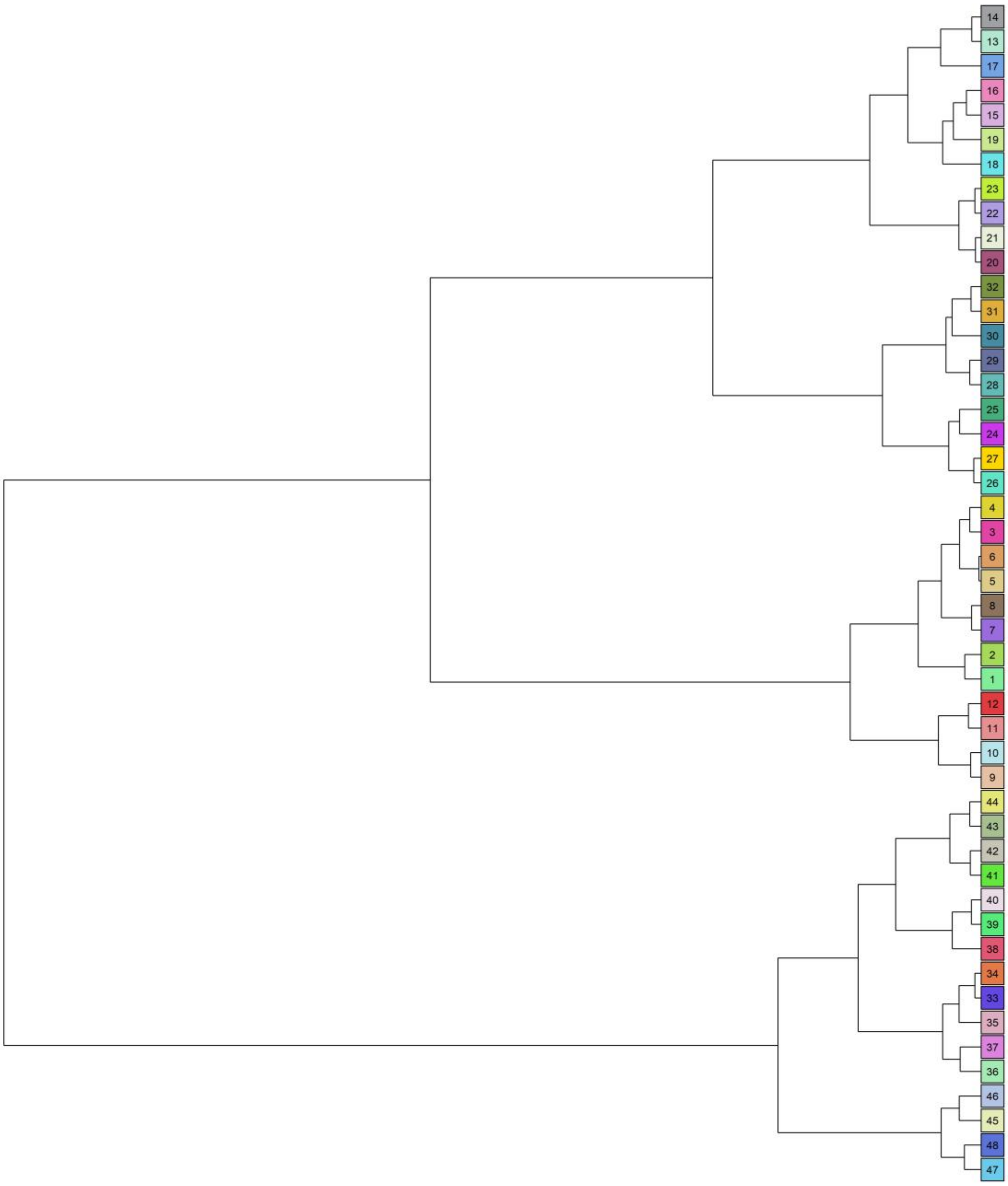


**Figure S8**. Environmental similarities among Seed Transfer Zones (STZs). Cluster analysis based on the mean values of the five Principal Component (PC) axes and scaled latitude and longitude.
